# The temporal dynamics of background selection in non-equilibrium populations

**DOI:** 10.1101/618389

**Authors:** Raul Torres, Markus G Stetter, Ryan D Hernandez, Jeffrey Ross-Ibarra

## Abstract

Neutral genetic diversity across the genome is determined by the complex interplay of mutation, demographic history, and natural selection. While the direct action of natural selection is limited to functional loci across the genome, its impact can have effects on nearby neutral loci due to genetic linkage. These effects of selection at linked sites, referred to as genetic hitchhiking and background selection (BGS), are pervasive across natural populations. However, only recently has there been a focus on the joint consequences of demography and selection at linked sites, and empirical studies have sometimes come to apparently contradictory conclusions as to their combined effects. In order to understand the relationship between demography and selection at linked sites, we conducted an extensive forward simulation study of BGS under a range of demographic models. We found that the relative levels of diversity in BGS and neutral regions vary over time and that the initial dynamics after a population size change are often in the opposite direction of the long-term expected trajectory. Our detailed observations of the temporal dynamics of neutral diversity in the context of selection at linked sites in non-equilibrium populations provides new intuition about why patterns of diversity under BGS vary through time in natural populations and help reconcile previously contradictory observations. Most notably, our results highlight that classical models of BGS are poorly suited for predicting diversity in non-equilibrium populations.

## Introduction

The effects of natural selection and demography on neutral genetic diversity within populations have long been of interest in evolutionary and population genetics. Recent efforts in sequencing tens of thousands of genomes across a multitude of species have yielded new and valuable insights into how these two forces of evolution have shaped extant patterns of genomic variation. Yet, while the theoretical underpinnings of the effects of natural selection and demography on genetic diversity have been investigated for decades (Maynard Smith and Haigh 1974; Nei *et al*. 1975; Maruyama and Fuerst 1984, 1985; Kaplan *et al*. 1989; Tajima 1989; Charlesworth *et al*. 1993; Hudson and Kaplan 1995; Nordborg et *al*. 1996), detailed investigation into how they jointly act to create patterns of diversity in different populations remains lacking.

Both theory and empirical observation have long shown that patterns of neutral genetic variation can vary regionally across the genome as a function of recombination rate (Maynard Smith and Haigh 1974; Begun and Aquadro 1992). This is because natural selection operating on selected sites not only decreases genetic variation at the focal site but can also lead to decreases in nearby neutral genetic diversity due to genetic linkage (Cutter and Payseur 2013). These effects, known as genetic hitchhiking (Maynard Smith and Haigh 1974) (in which neutral variants rise to high frequency with adaptive variants) and background selection (Charlesworth et *al*. 1993) (BGS; in which neutral variants are removed along with deleterious variants) can be widespread across the genome (Elyashiv et *al*. 2016). Evidence for selection at linked sites has been found across an array of species, including *Drosophila melanogaster* (Begun and Aquadro 1992; Charlesworth 1996; Andolfatto 2007; Sella *et al*. 2009; Comeron 2014; Elyashiv *et al*. 2016), mice (Keightley and Booker 2018), wild and domesticated rice (Flowers *et al*. 2011; Xu *et al*. 2012), *Capsella* (Williamson *et al*. 2014), monkeyflowers (Stankowski *et al*. 2018), maize (Beissinger *et al*. 2016), and humans (Sabeti *et al*. 2002; Reed *et al*. 2005; Voight *et al*. 2006; McVicker *et al*. 2009; Cai *et al*. 2009; Hernandez *et al*. 2011; Lohmueller *et al*. 2011).

Demographic change can also impact patterns of diversity across the genome. For example, neutral theory predicts that the amount of genetic diversity is proportional to a population’s effective population size (*N_e_*), such that changes in *N_e_* should result in concomitant changes to diversity (Kimura 1983). One of the most common forms of a population size change is a population bottleneck, whereby populations suffer a large decrease in size, often followed by an expansion. Some of the ways bottlenecks can occur include: domestication events (Doebley *et al*. 2006; Tang *et al*. 2010; Wiener and Wilkinson 2011; Gaut *et al*. 2018), seasonal or cyclical fluctuations in population size (Elton 1924; Ives 1970; Itoh et al. 2009; Norén and Angerbjörn 2014), and founder events (David and Capy 1988; Dlugosch and Parker 2008; Henn *et al*. 2012). Notably, while the rate of loss of diversity in response to a population contraction is quite fast, the recovery of diversity following a population increase can be slow (Charlesworth 2009). As a result, large contemporary populations may still exhibit patterns of low average genetic diversity if their population size was much smaller in the recent past. In humans, this is clearly evident in European and Asian populations due to the out-of-Africa bottleneck (Auton *et al*. 2015).

**add mention of pi and xi stats** Because selection at linked sites and demography are both pervasive forces across a multitude of species, the characterization of how these two forces interact with one another is necessary in order to develop a full picture of the determinants of neutral genetic diversity. The efficiency of natural selection scales proportionally with *N_e_* (Ohta 1973) and the impact of selection at linked sites on neutral diversity is likely to be greater in larger populations (Kaplan *et al*. 1989; Cutter and Payseur 2013; Corbett-Detig *et al*. 2015) (but see Gillespie (2001); Santiago and Caballero (2016)). Further, demographic changes can also increase (in the case of bottlenecks) or decrease (in the case of expansions) the rate of drift. It is therefore plausible that the rate at which diversity at a neutral locus is perturbed by selection at linked sites could be highly dependent on both the current as well as long-term *N_e_* of the population. This competition between the strength of selection at linked sites (which increases with the census size *N*) and genetic drift (which decreases with census N) may be a key contributor to the limited range of diversity observed among species despite much larger observed differences in census size (Lewontin 1974; Gillespie 2001; Leffler *et al*. 2012; Corbett-Detig *et al*. 2015; Santiago and Caballero 2016). However, selection at linked sites alone may not be sufficient to explain the discrepancy between observed diversity and census populations sizes (Coop 2016), and the action of both demography and selection at linked sites in concert may provide a better model. Moreover, the heterogeneous structure of selection at linked sites across the genome may yield different responses to demography and population splits through time (Burri 2017), and the resulting effects on patterns of differentiation and divergence also remain largely unexplored (but see *e.g*. Stankowski *et al*. 2019).

Many models of selection at linked sites were also formulated with the assumption that the population is large enough (or selection strong enough) such that mutation-selection balance is maintained (Charlesworth *et al*. 1993; Zeng 2013; Nicolaisen and Desai 2013). However, non-equilibrium demographic change may break such assumptions and forces other than selection may drive patterns of variation in regions experiencing selection at linked sites. For example, during the course of a population bottleneck, genetic drift may transiently dominate the effects of selection at many sites such that traditional models of selection will poorly predict patterns of genetic diversity. Additionally, in regions affected by selection at linked sites, the impact of genetic drift may be exacerbated because of lower *N_e_* in those regions, resulting in greater losses to diversity than expected by the action of demography alone. A recent review by Comeron (2017) included an initial investigation into the impact of demography on diversity in regions under BGS and suggested a dependency on demographic history. Recent empirical work in maize and humans has also demonstrated a strong interaction between demography and selection at linked sites (Beissinger *et al*. 2016; Torres *et al*. 2018). Yet these studies also demonstrate the need for a deeper understanding of the interaction between these forces, as they observe contrasting patterns of diversity in populations that have undergone a bottleneck and expansion.

In order to more fully explore the joint consequences of demography and selection at linked sites, in this study we conducted extensive simulations of different demographic models jointly with the effects of BGS. We find that the time span removed from demographic events is critical for populations experiencing non-equilibrium demography and can yield contrasting patterns of diversity that may reconcile apparently contradicting results (Beissinger *et al*. 2016; Torres *et al*. 2018). Additionally, the sensitivity of genetic diversity to demography is dependent on the frequency of the alleles being measured, with rare variants experiencing more rapid dynamic changes through time.

Our results demonstrate that traditional models of selection at linked sites may be poorly suited for predicting patterns of diversity for populations experiencing recent demographic change, and that the predicted forces of BGS become apparent only after populations begin to approach equilibrium. Importantly, even simple intuition about the effect of selection at linked sites may lead to erroneous conclusions if populations are assumed to be at equilibrium. These results should motivate further research into this area and support the use of models that incorporate the joint effects of both demography and selection at linked sites.

## Materials and Methods

### Simulation model

We simulated a diploid, randomly mating population using fwdpy11 v0.1.2a (*https://github.com/molpopgen/fwdpy11*), a Python package using the fwdpp library (Thornton 2014). Selection parameters for simulating BGS followed those of Torres *et al*. (2018), with deleterious variation occurring at 20% of sites across a 2 Mb locus and the selection coefficient, *s*, drawn from two distributions of fitness effects (DFEs). Specifically, 13% of sites were drawn from a gamma distribution (parameterize as shape *α* = 0.0415 and rate *β* = 80.11) and seven percent from a distribution with *α* = 0.184, *β* = 6.25. These distributions mimic the DFEs inferred across non-coding and coding sites within the human genome (Boyko *et al*. 2008; Torgerson *et al*. 2009). Fitness followed a purely additive model in which the fitness effect of an allele was 0, 0.5s, and *s* for homozygous ancestral, heterozygous, and homozygous derived genotypes, respectively. Per base pair mutation and recombination rates also followed those of Torres *et al*. (2018) and were 1.66 × 10^−8^ and 8.2 × 10^−10^, respectively. We included a 200 kb neutral locus directly flanking the 2 Mb deleterious locus in order to observe the effects of BGS on neutral diversity. For all simulations, we simulated a burn-in period for 10N generations with an initial population size of 20,000 individuals before simulating under 12 specific demographic models. The demographic models included one demographic model of a constant sized population (model 1) and eleven non-equilibrium demographic models incorporating bottlenecks and expansions (models 2-12; Table 1; Figures S1-S2). For each demographic model, we also conducted an identical set of neutral simulations without BGS by simulating only the 200 kb neutral locus. Each model scenario was simulated 5,000 times.

**Table 1.** Demographic parameters for models 1-12

| demography type | model | ancestral population size<br>($N [N_{anc}]$) | bottleneck/expansion population size<br>($N$) | bottleneck duration<br>($N_{anc}$ generations) | expansion duration<br>($N_{anc}$ generations)* | final population size<br>($N$) |
| --- | --- | --- | --- | --- | --- | --- |
| constant | model 1 | 20000 | NA | NA | NA | 20000 |
| bottleneck | model 2 | 20000 | 2000 | 1 | NA | 2000 |
|  | model 3 | 20000 | 400 | 1 | NA | 400 |
| expansion | model 4 | 20000 | 40000 | NA | 1 | 40000 |
| bottleneck-expansion<br>(ancient) | model 5 | 20000 | 2000 | 0 | 1 | 200000 |
|  | model 6 | 20000 | 400 | 0 | 1 | 200000 |
|  | model 7 | 20000 | 2000 | 0.05 | 0.95 | 200000 |
|  | model 8 | 20000 | 400 | 0.05 | 0.95 | 200000 |
| bottleneck-expansion<br>(recent) | model 9 | 20000 | 2000 | 0 | 0.1 | 200000 |
|  | model 10 | 20000 | 400 | 0 | 0.1 | 200000 |
|  | model 11 | 20000 | 2000 | 0.05 | 0.05 | 200000 |
|  | model 12 | 20000 | 400 | 0.05 | 0.05 | 200000 |

\*population expansion in models 5-12 is exponential, but in model 4 population expansion is instantaneous

### Diversity statistics and bootstrapping

After the burn-in period, we measured genetic diversity (*π*) and singleton density (*ξ* the number of singletons observed within a window) within 10 kb windows across the 200 kb neutral locus every 50 generations using a random sample of 400 chromosomes. We measured *π* and *ξ* for each demographic model by taking the mean of these values across each set of 5,000 replicate simulations. For neutral simulations, we annotated *π* and *ξ* as *π*_0_ and *ξ_0_*, respectively. We took the ratio of these statistics (i.e., *π*/*π*_0_ and *ξ*/*ξ*_0_) in order to measure the relative impact of BGS within each demographic model. We bootstrapped the diversity statistics by sampling with replacement the 5,000 simulated replicates of each demographic model to generate a new set of 5,000 simulations, taking the mean of *π* and *ξ* across each new bootstrapped set. We conducted 10,000 bootstrap iterations and generated confidence intervals from the middle 95% of the resulting bootstrapped distribution.

### Calculations of expected BGS

To calculate the predicted equilibrium *π*/*π*_0_, we first used equation 14 of Nordborg *et al*. (1996), but modified it to incorporate two gamma distributions of fitness effects. Additionally, in order to properly model our simulations, we only calculated the effects of BGS on one side of the selected locus. This resulted in the following modified equation:

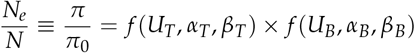

The first term of the equation (*f*(*U_T_*, *α_T_*, *β_T_*)) models the effects of BGS due to selection on non-coding sites according to the gamma DFE inferred by Torgerson *et al*. (2009), and the second term of the equation (*f* (*U_B_*, *α_B_*, *β_B_*)) models the effects of BGS due to selection on coding sites according to the gamma DFE inferred by Boyko *et al*. (2008). Each of these is modeled following:

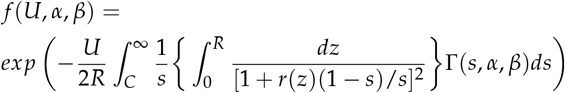

Here, *R* is the total length of the selected locus in bp, *U* is the total deleterious mutation rate across the selected locus, *r*(*z*) is the genetic map distance between a neutral site and a deleterious mutation, and *s* is the selection coefficient of a deleterious mutation.

Because *N* is not explicitly included in this model of BGS, we followed previous work (Charlesworth 2012; Comeron 2014) in truncating selection at some value *C* (represented in the integral 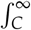). Here, *C* represents the minimum selection coefficient (s) that is treated as deleterious for the model. This step effectively excludes neutral mutations from the model that should not contribute to BGS, and can be modulated to mimic small or large populations (by increasing or decreasing *C*, respectively). This truncation step also affects the values used for *U* in the above equation, resulting in specific values of *U* for each DFE. We simulated different population sizes to equilibrium under our BGS simulation model to see how well the modified version of the classic model fit populations of different *N* for different values of *C* (Figure S3). Despite the fact that our simulations potentially break assumptions of the model (e.g., mutant alleles at frequencies rare enough that higher-order terms can be ignored and multiplicative fitness effects across loci), we observed a generally good fit of our resulting observed *π*/ *π*_0_ to the expectations of Nordborg *et al*. (1996) (Figure S3).

Because no single value of *C* provided an estimate of BGS that was robust to the population sizes simulated in our demographic models, we also fit a log-linear model to the observed values of *π*/ *π*_0_ for varying sizes of *N* (Figure S3). The resulting best fit model to the observed data was:

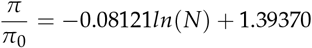

For each generation of our demographic models, we calculated the long-term effective population size (*N_e_*) by applying the following equation:

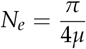

Here, we substituted the mutation rate used in our simulations for *μ* (1.66 × 10^−8^). For *π*, we used the mean observed per-site diversity across the 200 kb neutral region from each set of 5,000 neutral simulations (i.e., *π*_0_).

Using the fitted log-linear model of population size and *π*/*π*_0_ and the calculations of long-term *N_e_* described above, we also estimated *π*/*π*_0_ for each generation in our demographic models by substituting the estimated long-term *N_e_*, for *N* in −0.08121*ln*(*N*) + 1.39370.

Simulation and analysis code are available at https://github.com/RILAB/BGS_sims/.

## Results

### Background selection under instantaneous population size change

We first present the joint effects of demography and BGS under simple demographic models with a single instantaneous change in size (models 2-4; Figure S1). While our simulations incorporated a 200 kb neutral region, we first focused on patterns of diversity generated within the 10 kb window nearest to the 2 Mb locus experiencing purifying selection, as this is where BGS is strongest. Doing so allowed us to observe any change in the dynamics of *π* and *ξ* as they approached new population equilibria resulting from a change in size. In the simple bottleneck models (models 2-3) we observed the expected strong decrease in *ξ* and *π* following population contraction in models of both BGS and neutrality (Figure 1). Similarly, we observed the expected rapid increase in *ξ* compared to *π* in our model of a simple population expansion (model 4; Figure 1). In all cases, values of *ξ* and *π* were lower in models with BGS and changed more quickly relative to their initial value than in the neutral case (Figure S4).

**Figure 1.**
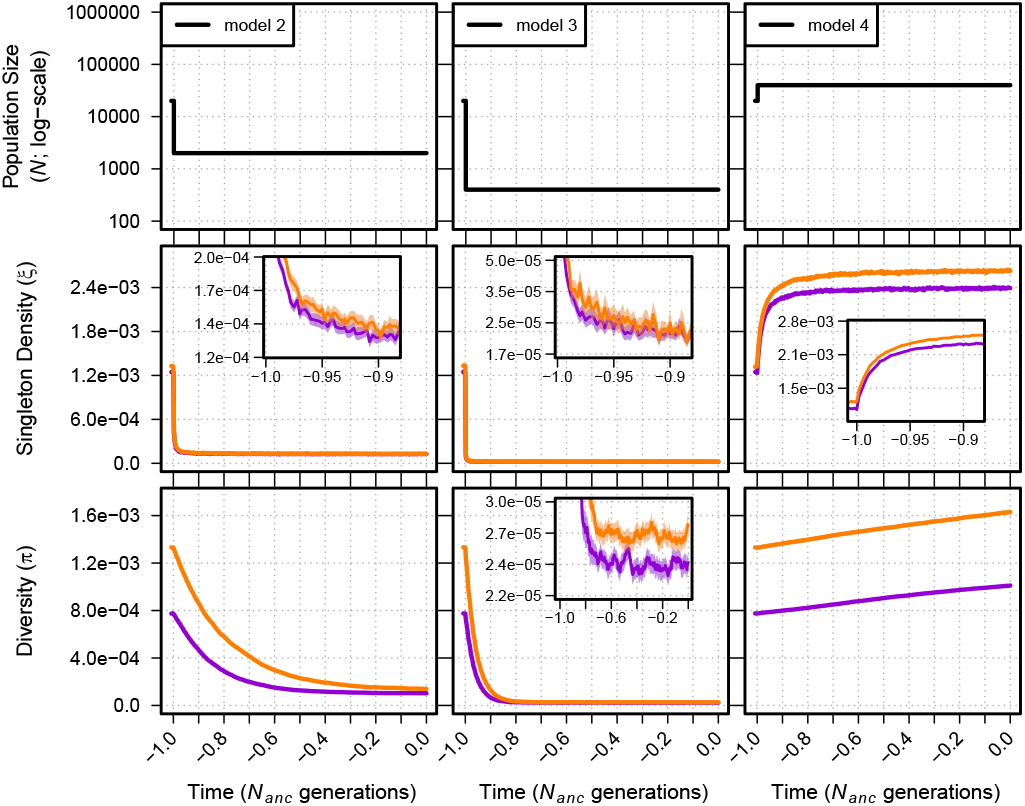
Singleton density (*ξ* per site) and diversity (*π* per site) for models 2-4. The top panel shows each demographic model; time proceeds forward from left to right and is scaled by the *N* of the population at the initial generation (*N_anc_*; 20,000 individuals). Diversity statistics are shown for neutral simulations (orange lines) and simulations with BGS (violet lines). Insets show diversity using a log scale for detail. Envelopes are 95% CIs calculated from 10,000 bootstraps of the original simulation data.

To examine the interaction of demography and selection observed in empirical data (Beissinger *et al*. 2016; Torres *et al*. 2018), we normalize *π* and *ξ* in models of BGS by their equivalent statistics generated under the same demographic model in the absence of any selection (*π*_0_ and *ξ*_0_). We observed that *π*/*π*_0_ and *ξ* / *ξ*_0_ were dynamic through time in response to demography, with changes occurring to both their magnitude and direction (Figure 2). Moreover, changes to *ξ*/*ξ*_0_ occurred more rapidly through time compared to *π*/*π*_0_. For example, in model 2 we observed a dip and rise in the *ξ*/*ξ*_0_ statistic relative to equilibrium (model 1) within the first ≈ 0.1 *N_anc_* generations (*N_anc_* refers to the size of the ancestral population prior to any demographic change). Yet, for the same model, *π*/*π*_0_ remained depressed for over 0.5 *N_anc_* generations (Figure 2). Similar patterns were observed for model 3, which experienced a greater reduction in size, although the pattern is less clear because of the greater sampling variance of *ξ*/*ξ*_0_ due to the overall lower number of singletons. In both population contraction models, *π*/*π*_0_ and *ξ*/ *ξ*_0_ appeared to plateau at levels above that of the equilibrium model (model 1). In contrast, we observed markedly different dynamics in our model of a simple population expansion (model 4). This included a sustained increase in *π*/*π*_0_ but only a transient increase in *ξ*/*ξ*_0_ which drops below the equilibrium model within the first ≈ 0.1 *N_anc_* generations.

**Figure 2.**
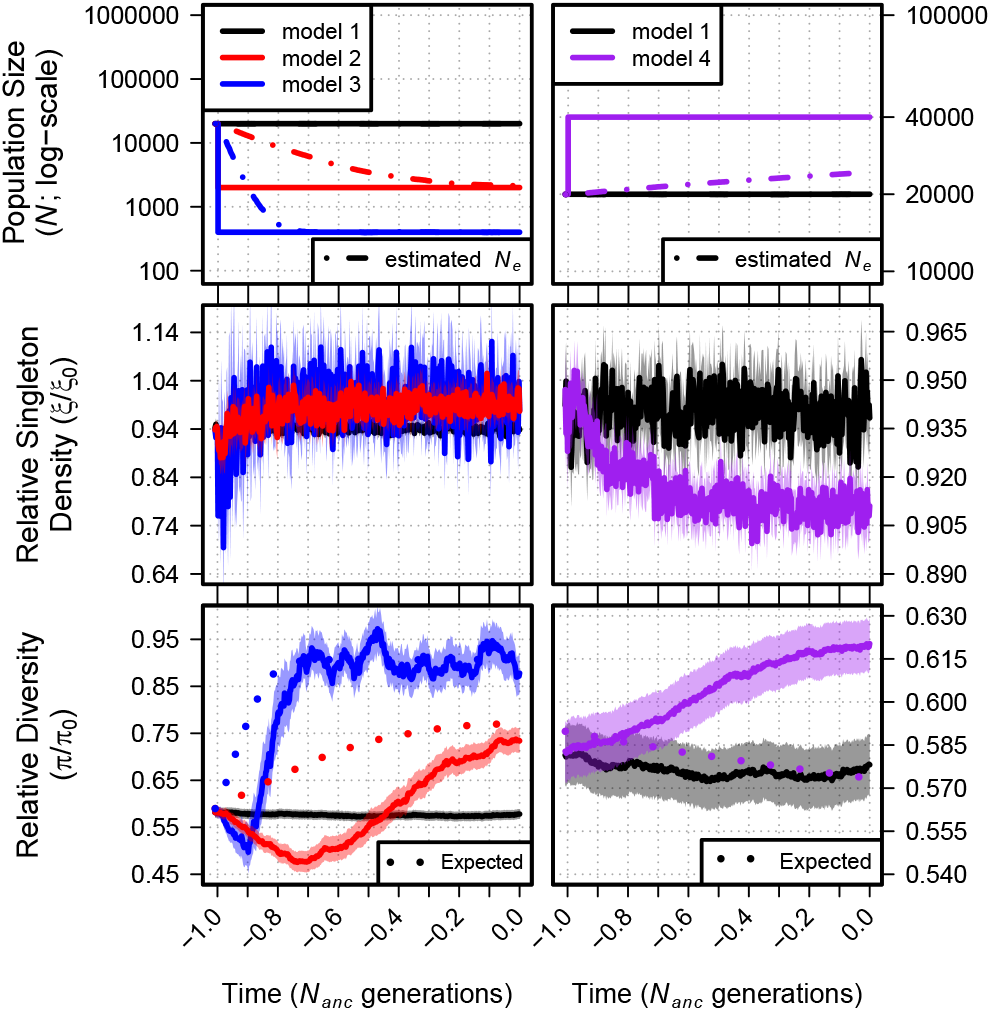
Relative singleton density (*ξ*/*ξ*_0_) and relative diversity (*π*/*π*_0_) across time for demographic models 1-4. The top panel shows each demographic model as in Figure 1. Black lines show *ξ*/*ξ*_0_ and *π*/*π*_0_ from simulations of a constant sized population (model 1). Dot-dashed lines in the top panels show the estimated *N_e_* from observed *π*_0_. Dotted lines in the bottom panel show the equilibrium expectation of *π*/*π*_0_ from a log-linear model of simulated BGS with the specific selection parameters and the estimated *N_e_* at each time point (see Figure S3). Envelopes are 95% CIs calculated from 10,000 bootstraps of the original simulation data.

Changes in population size should lead to changes in the rate of genetic drift and the efficacy of natural selection and, thus, changes in the magnitude of BGS over time. Indeed, under equilibrium conditions (and if mutations that are effectively neutral can be ignored) the classic model of BGS (Nordborg *et al*. 1996) predicts weaker BGS (with higher *π*/*π*_0_) for smaller populations and stronger BGS (with lower *π*/*π*_0_) for larger populations (Figure S3). To compare these predictions to those of our simple demographic models, at each generation for each model, we calculated *π*/*π*_0_ using a log-linear model fit to predict *π*/*π*_0_ from *N* (see Materials and Methods). In all three simple demographic models, we observed that changes in *π*/*π*_0_ over the short term differed qualitatively from the predicted *π*/*π*_0_ of the log-linear model (Figure 2; bottom panel). While the log-linear model predicts a higher value for *π*/*π*_0_ in a smaller population, we observed a transient drop in *π*/*π*_0_ directly after a contraction (models 2 and 3). Similarly, the log-linear model predicts a decrease in *π*/*π*_0_ in larger populations, but we instead observed an increase in *π*/*π*_0_ with a population expansion (model 4). The trajectory of *π*/*π*_0_ changed in our bottleneck models, eventually approaching the higher values predicted by the log-linear model and in line with the overall predictions of the classic model of BGS. In contrast, *π*/*π*_0_ in the expansion model continued to increase over the entire course of the simulation. To test if and when *π*/*π*_0_ for the expansion model reaches the lower value predicted by the log-linear model, we ran a limited set of simulations (2,000 total) for 11 *N_anc_* generations. We found that, indeed, *π*/*π*_0_ plateaued and then decreased relative to its starting value, eventually approaching the prediction of the log-linear model after ≈ 10 *N_anc_* generations (Figure S10). This was because, in the absence of selection, *π* plateaus more slowly when compared to *π* under BGS (Figure S11). Only once *π* under neutrality began to approach equilibrium did we begin to observe the prediction of the log-linear model.

In order to test whether stronger or weaker effects of selection change the overall patterns we observed with our simulations using a mixture of two DFEs, we also conducted simulations with *s* drawn from a point distribution (*γ* = *2Ns* = {0.1, 0.5, 2, 5, 10, 50, 100}) for models 3 and 4. The results displayed broadly similar patterns but with differing degrees of change in *π*/*π*_0_ or *ξ*/*ξ*_0_ depending on the strength of selection (Figure S12). For model 3, *π*/*π*_0_ increased as before as the population approached equilibrium, stabilizing near 1 except under models with the strongest selection. However, the transient changes in *π*/*π*_0_ seen in Figure 2 directly after population size change were much less evident in these simulations and were essentially absent in models with stronger selection. For *ξ*/*ξ*_0_, patterns were both more dynamic and more closely matched to those of Figure 2, with rapid transient decreases and increases occurring shortly after a contraction (model 3) or expansion (model 4), respectively.

### Background selection under bottleneck-expansion models

We built upon the simple two epoch demographic models to test more complex scenarios and better understand the relative effects of different events on patterns of diversity under BGS. Specifically, we simulated a population undergoing a contraction similar in size to models 2 and 3, but with a subsequent expansion to 400,000 individuals by the final generation (Figure S2; Table 1). These bottleneck-expansion models included both ancient (1.0 *N_anc_* generations in the past; models 5-8) and recent (0.1 *N_anc_* generations in the past; models 9-12) instantaneous bottlenecks with either an immediate return to growth (models 5-6, 9-10) or a sustained bottleneck (models 7-8, 11-12).

These models recapitulated several patterns observed in our simple bottleneck models, but with added dynamics. In all cases, diversity in models with BGS was both lower (Figures S5–S6) and changed more rapidly (Figures S7–S8) than in neutral simulations. Changes in diversity also occurred more quickly in models with a stronger or sustained bottleneck, and *ξ* again exhibited more rapid dynamics than did *π*. Mirroring results from our simple bottleneck scenarios, models with an ancient bottleneck (models 5-8) showed a transient decrease in *ξ*/*ξ*_0_ and *π*/*π*_0_ followed by an increase to higher values (Figure 3). Again, changes in *π*/*π*_0_ contrast with the expectations of the log-linear and classic models, where BGS is expected to become more efficient in larger populations resulting in an expected decrease in *π*/*π*_0_ through time (Figure 3, dotted lines). But while both *π*/*π*_0_ and /0 remain elevated in our simple bottleneck models, *ξ*/*ξ*_0_ in the bottleneck-expansion models shifts direction during the course of the expansion and begins to decline, eventually reaching values below that of the equilibrium population. Finally, because of the added complexity of the expansion following the population bottleneck, it is also likely that the increase in *π*/*π*_0_ for these models later in their demographic histories is also recapitulating the similar dynamics witnessed for model 4.

**Figure 3.**
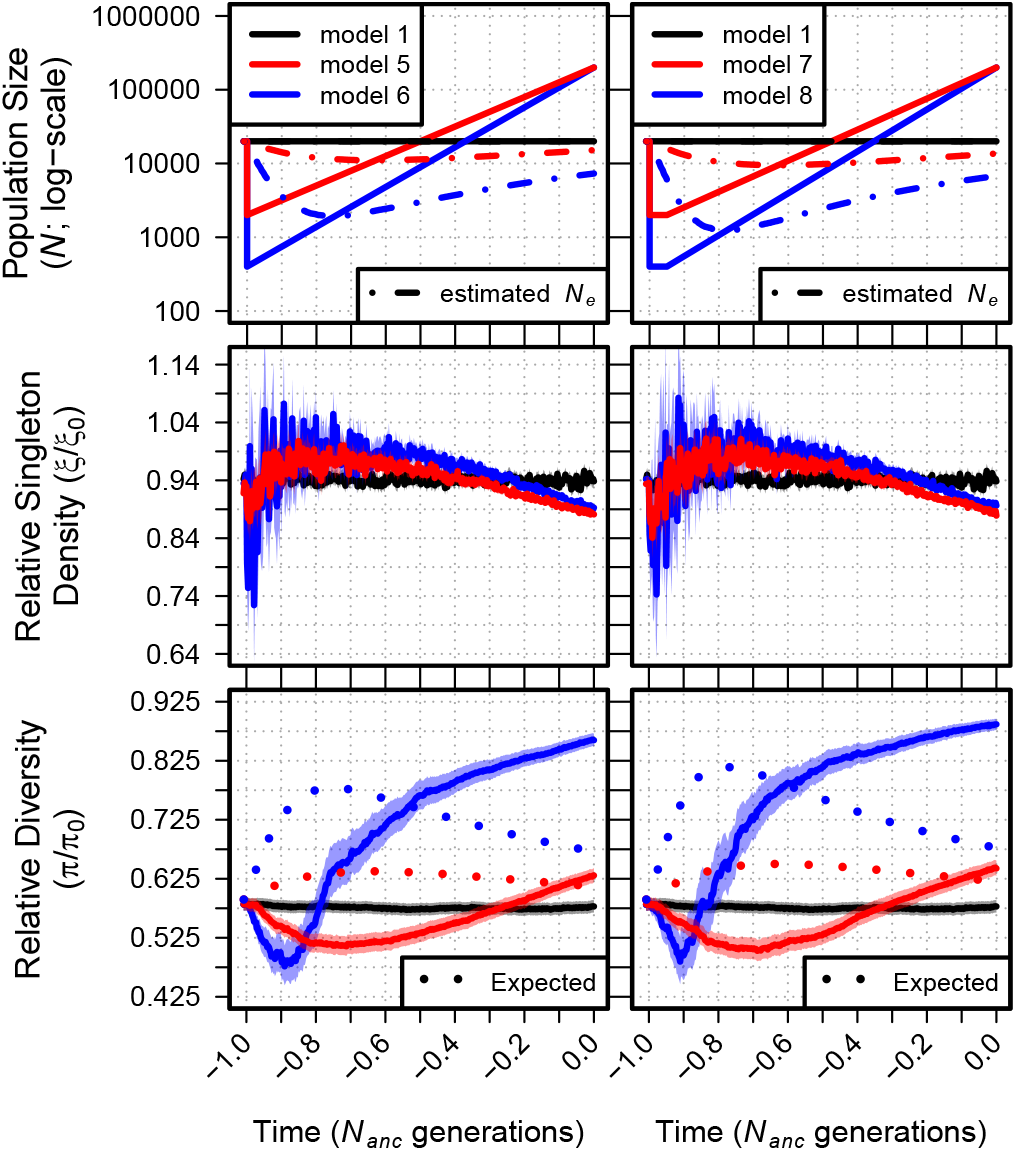
Relative singleton density (*ξ*/*ξ*_0_) and relative diversity (*π*/*π*_0_) across time for demographic models 1 and 5-8. The top panel shows each demographic model; time proceeds forward from left to right and is scaled by the *N* of the population at the initial generation (*N_anc_;* 20,000 individuals). Black lines show *ξ*/*ξ*_0_ and *π*/*π*_0_ from simulations of a constant sized population (model 1). Dot-dashed lines in the top panels show the estimated *N_e_* from observed *π*_0_. Dotted lines in the bottom panel show the equilibrium expectation of *π*/*π*_0_ from a log-linear model of simulated BGS with the specific selection parameters and the estimated *N_e_* at each time point (see Figure S3). Envelopes are 95% CIs calculated from 10,000 bootstraps of the original simulation data.

Though the trajectories of *π*/*π*_0_ and *ξ*/*ξ*_0_ were truncated for models in which the bottleneck occurred in the recent past (models 9-12; 0.1 *N_anc_* generations), they nonetheless appeared to behave qualitatively similar to ancient bottleneck models (Figure S9). It is worth noting, however, that because of the difference in timescale, the ending values of *ξ*/*ξ*_0_ and *π*/*π*_0_ from recent bottleneck models is in the opposite direction relative to model 1 when compared to models with longer demographic histories (models 5-8).

### Patterns of diversity across the 200 kb neutral region

We also measured patterns of *π*/*π*_0_ across time for the entire 200 kb neutral region. Doing so showed the characteristic “trough” structure of increasing relative diversity as a function of genetic distance from the deleterious locus (model 5 is shown in Figure 4, see Figure S13 for all models). Change in *π*/*π*_0_ over time generally followed patterns observed in the neutral window closest to the selected region. In all of our ancient bottleneck models (models 2-3, 5-8), for example, we see a decline in *π*/*π*_0_ across the entire region followed by an increase to levels higher than in the ancestral population. For recent bottlenecks (models 9-12) we see a consistent decline with no recovery and in our simple expansion model (model 4), *π*/*π*_0_ increases monotonically through time.

Yet, these general patterns obscure more subtle changes in the slope of *π*/*π*_0_ in the trough structure. In models with a stronger bottleneck (models 3, 6 and 8), where we expect the efficacy of selection to be most affected in the long term, we see that the slope *π*/*π*_0_ flattens over time, completely erasing the trough of diversity in the most extreme case without a recovery (model 3).

Finally, while *ξ*/*ξ*_0_ across the region largely followed patterns seen in the neutral window most proximal to the selected locus, a closer look across the 200 kb regions of most models yielded no clear patterns. Troughs were slightly apparent for the final generations of some models (models 5 and 7), but the stochasticity among 10 kb windows for *ξ*/*ξ*_0_ swamped any other patterns that might otherwise be evident.

**Figure 4.**
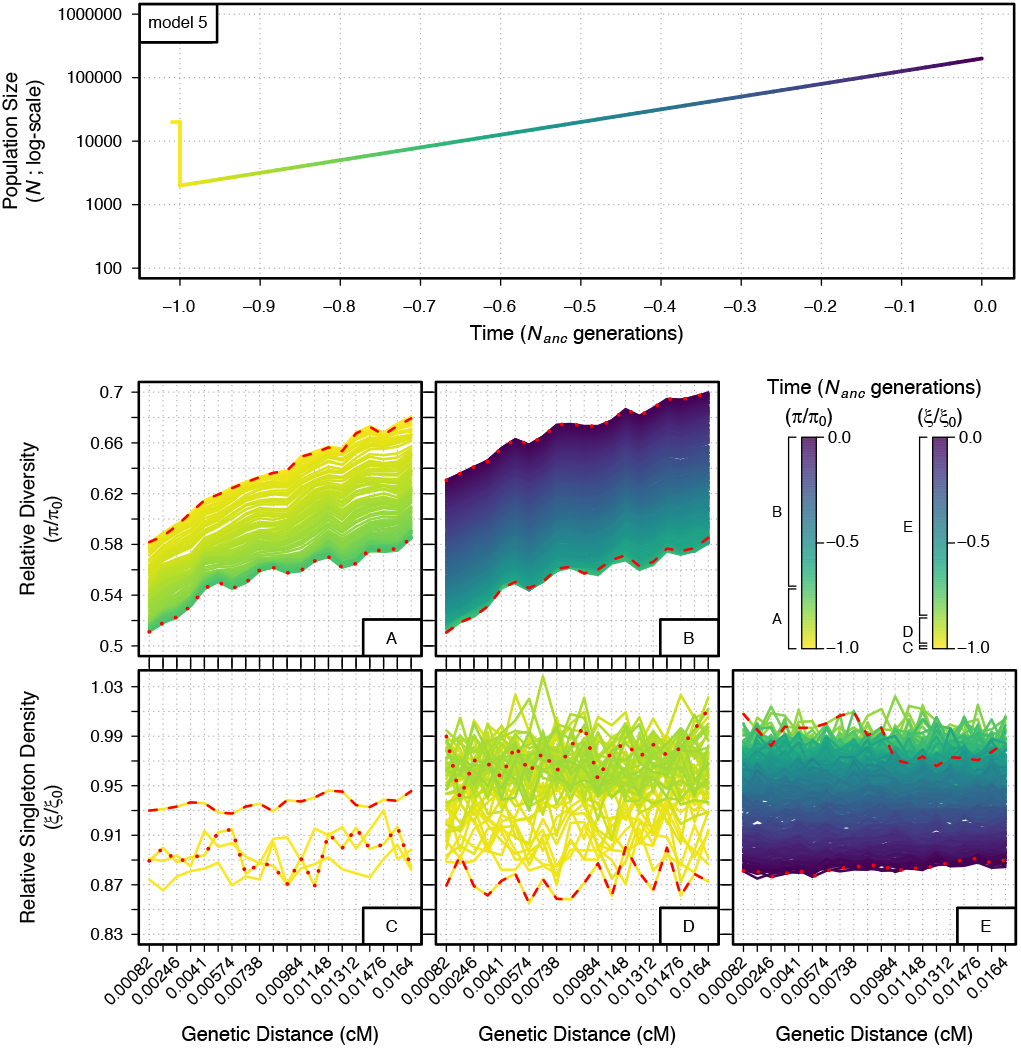
Temporal and spatial dynamics of relative diversity (*π*/*π*_0_) and singleton density (*ξ*/*ξ*_0_) under a bottleneck with expansion (model 5) across a neutral 200 kb region. The genetic distance of each 10 kb bin from the selected locus is indicated on the x-axis of the bottom panels. Each line measuring *π*/*π*_0_ and *ξ*/*ξ*_0_ in the bottom panels represents one of the 401 discrete generations sampled from the demographic model; colors follow the demographic model in the top panel (time is scaled as in Figures 1–3) and in the figure legend. Multiple plots are given in order to prevent overlap of the measurements between generations. Panels A and B show *π*/*π*_0_ through time from −1.0 to −0.71 *N_anc_* (A) and −0.71 to 0.0 *N_anc_* (B) generations. Panels C, D and E show / _0_ through time from −1.0 to −0.99 *N_anc_* (C), −0.99 to −0.85 *N_anc_* (D) and −0.85 to 0.0 *N_anc_* (E) generations. Red dashed lines and red dotted lines indicate the first and last generation measured within each plot, respectively.

## Discussion

### General patterns of diversity

A long history of both theoretical (Nei *et al*. 1975; Maruyama and Fuerst 1984, 1985) and empirical (Begun and Aquadro 1992; Cavalli-Sforza *et al*. 1994; Eyre-Walker *et al*. 1998) population genetics work has provided a clear picture of the impacts of demographic change on patterns of diversity in the genome. We know, for example, the impact of simple bottleneck and growth models on the allele frequency spectrum (Tajima 1989; Slatkin and Hudson 1991; Griffiths and Tavare 1994). Theory also offers clear direction on the long-term effects of decreases in effective population size on the efficacy of natural selection (Ohta 1973; Kimura 1983). Likewise, classical theory on background selection provides a solid expectation for the effects of selection at linked sites on diversity in populations at demographic equilibrium (Nordborg *et al*. 1996). For instance, the reduction in genetic diversity under the influence of BGS increases with increasing population size in equilibrium populations (Nordborg *et al*. 1996, Figure S3).

Despite these efforts, there have been surprisingly few investigations addressing the expected patterns from the interaction of demography and selection at linked sites in the context of BGS (Zeng 2013; Nicolaisen and Desai 2013; Ewing and Jensen 2016; Comeron 2017; Rettelbach *et al*. 2019). There also remains substantial confusion in empirical population genetic analyses, with authors often equating long-term predictions of change in effective population size on the efficacy of natural selection to short-term responses under non-equilibrium demography (Brandvain and Wright 2016). Here, we use simulations and analysis of different demographic models with and without BGS to show that predictions from such equilibrium models generally fail to hold up over shorter time scales. We find that the predicted impacts of the combined effects of demography and selection at linked sites depend strongly on the details of the demographic model as well as the timing of sampling.

In each of our models, the initial effects observed are driven primarily by the stochastic effects of drift. For example, the loss of diversity in the first few generations after a population decline occurs equally across the entire region, independent of the distance from the selected region (Figure S13). These effects occur more rapidly in regions undergoing BGS than neutral regions (Figure 5). For example, while equilibrium models predict that the effects of BGS should be attenuated in populations with lower *N_e_* due to the decreased efficacy of purifying selection, we instead observed a drop in *π*/*π*_0_ after the bottleneck and a more rapid decrease in models with a stronger bottleneck (Figures 2 and 3). Similarly, while theory predicts a decrease in *π*/*π*_0_ in larger populations, we instead observed that the initial response after population expansion was an increase in *π*/*π*_0_ (Figure 2). These observations make it clear that the combined effects of demography and BGS on *π*/*π*_0_ immediately following a reduction in *N_e_* are not driven by a change in the efficacy of natural selection, but rather by changes to the rate of drift in regions undergoing BGS.

**Figure 5.**
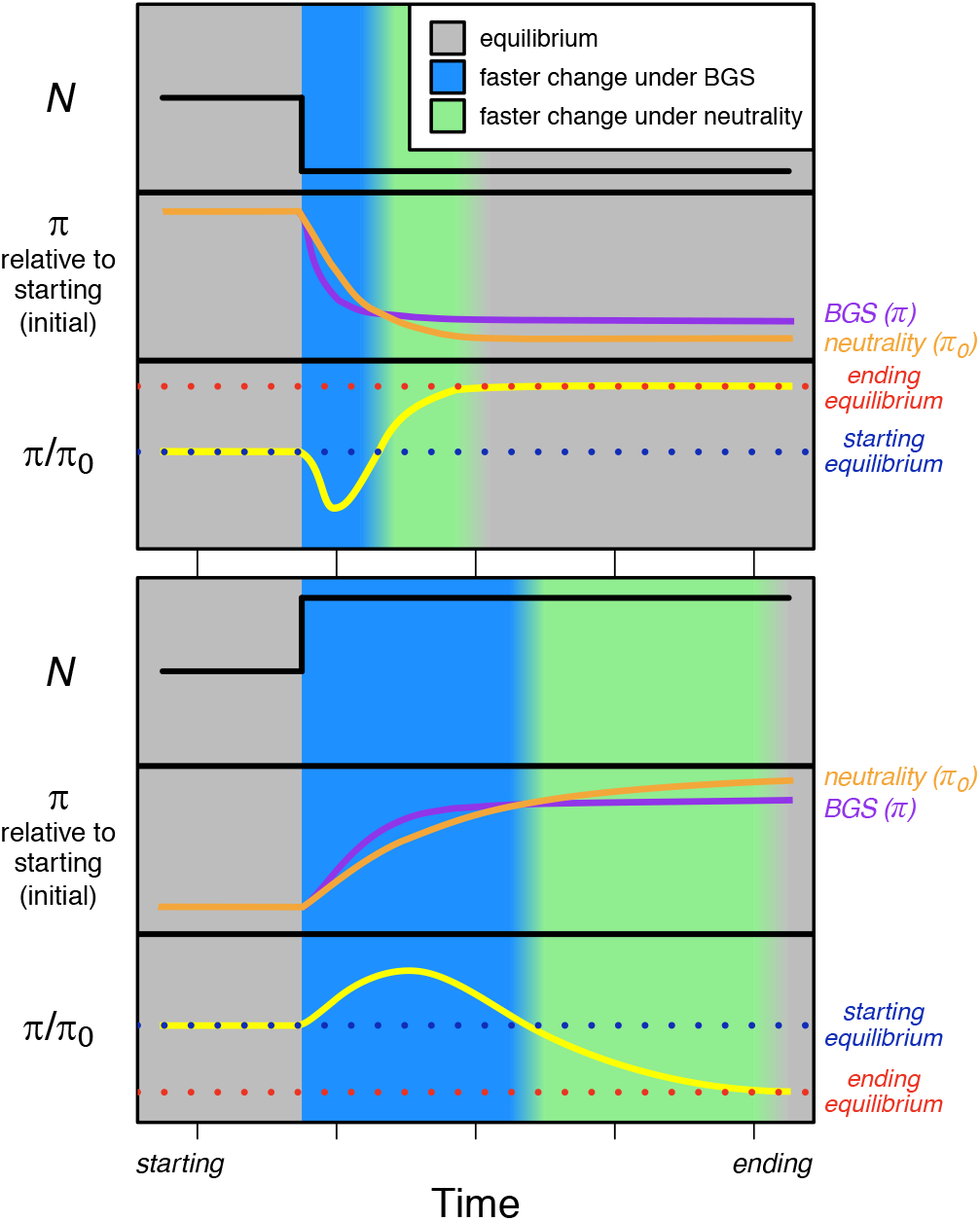
Schematic of the temporal dynamics of diversity under neutrality (*π*_0_; orange lines) and BGS (*π*_0_ violet lines) for a demographic bottleneck and an expansion. Relative diversity (*π*/*π*_0_; yellow lines) is shown in the bottom section of each figure panel with equilibrium points before demographic change (blue dotted lines) and after demographic change (red dotted lines) shown. Background colors represent epochs of time where the change in diversity is faster under BGS (blue) or neutrality (green).

Although the initial changes in diversity are dominated by the impacts of demography, as population size shifts, the efficacy of natural selection begins to change as well. In our simple bottleneck models, *π*/*π*_0_ stops declining and begins to increase, eventually reaching higher values as expected under equilibrium (Figure 2). This change reflects the inability of a smaller population to select against new deleterious mutations, rendering these alleles effectively neutral and decreasing the effects of BGS. These effects are countered in larger, growing populations, which is presumably why we see the rate of increase of *π*/*π*_0_ slow and eventually plateau in models incorporating both bottlenecks and growth (Figure 3). In addition, the rate of change in *π*/*π*_0_ is also diminished, with slower approaches to equilibrium taking place as a function of larger population size. Indeed, in our simple expansion models, patterns of diversity approach equilibrium expectations only after approximately ≈ 10*N_anc_* generations (Figure S10).

Changes in the efficacy of selection are also readily observed in comparisons of relative diversity across windows varying in recombination distance from the selected region (Figure SI 3). Diversity in the ancestral population increases with distance from the selected regions as expected under classical models of BGS at equilibrium (Nordborg *et al*. 1996) and observed in previous studies (Hernandez *et al*. 2011; Beissinger *et al*. 2016). But while the slope of this relationship remains constant in the generations initially following population size change, for our simple bottleneck models, it begins to flatten through time, reflecting a lowered effective population size and a concomitant weakened efficacy of natural selection (Figure S13; models 2-3).

The diversity-reducing effects of BGS have often been modeled as a reduction in *N_e_* (Charlesworth *et al*. 1993), though we caution that the effects of BGS on the SFS cannot be simplified to this extent (Cvijović *et al*. 2018). Like a reduction in *N_e_,* however, BGS exacerbates the stochastic process of drift. Because the relevant timescale for allele frequency evolution is scaled by the rate of drift (Crow and Kimura 1970), both the reduction and recovery of *π*/*π*_0_ to equilibrium levels happen over fewer generations in populations with stronger bottlenecks and in regions impacted by BGS. We see this borne out in comparisons of models with stronger (Figure 2) or more sustained (Figure 3) bottlenecks, as well as comparisons of models with BGS to their equivalent neutral scenarios (Figures S4, S7, S8, and S11). This differential scaling also contributes to the observed lag in time in reaching equilibrium for neutrality relative to BGS (Figure 5) and the slower rate of change observed in expanding populations (Figure 3), as increases in the effective size attenuate the rate of drift.

The timing and magnitude of changes in diversity also depend on the range of allele frequencies assayed. We demonstrate this by analyzing changes in singleton density (*ξ*) along with overall patterns of nucleotide diversity. Because singleton variants represent very recent mutations, changes in *ξ*/*¾*_0_ respond more quickly to changes in *N*. In our simple expansion model, for example, while *π*/*π*_0_ increases for ≈ 2*N_anc_* generations (Figure S10), we see a relatively rapid increase in *ξ*/*¾*_0_ followed by a decrease as the larger population size increases the efficacy of selection against new deleterious mutants. And while theoretical predictions for *ξ*/*ξ*_0_ are not as straightforward (because of the dependency of distortions to the site-frequency spectrum on sample size (Cvijović *et al*. 2018)), singleton density in the simple expansion model quickly stabilizes at a new value below that of the ancestral population, consistent with having reached a new equilibrium value. However, signals using rare frequency bins such as are inherently more difficult to capture, partly because they are less affected than *π* since BGS perturbs common frequency bins of the SFS more than rare ones (Cvijovicć *et al*. 2018). In addition, we observe much higher variance for *ξ*/*ξ*_0_ compared to *π*/*π*_0_.

Finally, although we have simulated only one complex DFE under a mixture distribution of selection coefficients for new mutations (see Materials and Methods), this distribution will also play an important role in determining the threshold above which new mutations contribute to the effects of linked selection. For example, while our DFE had a mean of *2N_anc_s* = 424, it is also characterized by an extremely long tail and ≈ 75% of deleterious mutations will only have a *s* ≤ 10^−3^, which is equivalent to *2N_anc_s* = 40. These features add additional complexity to both the initial and long-term dynamics of diversity after demographic change when compared to simulations of BGS using a single value of *s* (Figure S12). In simulations using our wide DFE, the transient drop in *π*/*π*_0_ following a contraction was stronger than in simulations using a single *s*, but the long-term qualitative results differed as well: *π*/*π*_0_ was initially higher than models where *2N_anc_s* = {2, 5, 10} but is lower after the population reaches its new equilibrium. It is thus clear that the details of both the short-and long-term changes in diversity as a result of the interaction of demography and selection will depend on features of the DFE as well.

### Conflicting signals in maize and humans

One of the motivations for the work presented here is the fact that empirical analyses evaluating the impact of demography on selection at linked sites have come to conflicting conclusions (Torres *et al*. 2018; Beissinger *et al*. 2016). Beissinger *et al*. (2016) compared domesticated maize to its wild ancestor teosinte, finding higher *π*/*π*_0_ but lower *ξ*/*ξ*_0_. In a similar analysis in humans, Torres *et al*. (2018) found lower *π*/*π*_0_ but higher *ξ*/*ξ*_0_ in non-African compared to African populations.

The fact that both maize and non-African human populations have undergone a population bottleneck and expansion over a similar timescale (on the order of ≈ *0.1N_anc_* generations) makes the contrasting results from these papers initially somewhat surprising. But despite their qualitatively similar demographies, there are a number of factors that complicate direct comparison between humans and maize and highlight difficulties in inferring the action of linked selection in empirical data. For example, while the domestication process in maize is widely thought to have led to a population bottleneck and subsequent expansion (Eyre-Walker *et al*. 1998; Tenaillon *et al*. 2004; Wright *et al*. 2005; Beissinger *et al*. 2016; Wang *et al*. 2017; Bellon *et al*. 2018), there is little agreement on the magnitude and timing of these effects, and estimates of the modern maize population vary by several orders of magnitude. And while the estimated demography of African populations is relatively stable compared to that of non-Africans (Torres *et al*. 2018), the demography of teosinte is not well understood but is likely to include substantial nonequilibrium dynamics (Wang *et al*. 2017). The distribution of fitness likely differs between the species as well. Compared to the mixture of two DFEs used by Torres *et al*. (2018), the best-fit gamma distribution for the only published DFE estimated for maize exhibits a nearly 10-fold higher mean (Pophaly and Tellier 2015). Finally, the distribution of functional sites in the genome differs between humans and maize – genes are longer in humans, leading to smaller intergenic spaces and perhaps shorter average distances to the nearest functional site.

Nonetheless, simulations combining demography and background selection highlight plausible scenarios that could result in the differences seen between maize and humans. In models 9-10 (Figure S9), for example, patterns of diversity ≈ 0.1*N_anc_* generations after the bottleneck qualitatively match those of Torres *et al*. (2018), with *π*/*π*_0_ lower and *ξ*/*ξ*_0_ higher than seen in the ancestral population. In their simulations of a genic region using a single s, Beissinger *et al*. (2016) found no differences in *π*/*π*_0_ between maize and teosinte but much lower *ξ*/*ξ*_0_, providing some evidence to support their observed findings. As demographic inferences comes with uncertainty, it is possible the population bottleneck during maize domestication was potentially weaker or the expansion began much sooner. In such a case, something closer to our model 4 (Figure 2) might be a reasonable comparison. Indeed, in that scenario, 0.1*N_anc_* generations after the expansion we see *π*/*π*_0_ has increased and *ξ*/*ξ*_0_ decreased compared to the ancestral population, similar to the observations of Beissinger *et al*. (2016). Improved sampling and more careful modeling of both demography and the DFE is likely required, however, to demonstrate whether the observed results in maize can be entirely explained by the interaction of BGS and demography.

### Implications for empirical data

Combined with our simulation results, the difficulty of interpreting what appear as straightforward differences between maize and humans suggests that inferences about linked selection from empirical data is likely to be difficult without careful consideration of demography. Indeed, even the relationship between *π* and recombination changes over time in our models (Figure S13), highlighting the importance of incorporating demography into models that use such information to make inference about linked selection. Most of the work to date using either theory (e.g., Elyashiv *et al*. 2016; Corbett-Detig *et al*. 2015; Rettelbach *et al*. 2019) or simulation (Stankowski *et al*. 2019) makes use of classic equations that assume populations are at equilibrium.

Given these complexities, what considerations should researchers interested in empirical analysis keep in mind? While simulation results suggest that BGS is unlikely to strongly affect the ability to detect outliers via selection scans using *F_ST_* (Matthey-Doret and Whitlock 2019), we argue here against using simple approximations based on equilibrium models to infer the relative importance of demography and selection in patterning diversity along the genome. For researchers interested in assessing the impacts of demography and linked selection, we first recommend careful consideration of how *π*_0_ is estimated, making efforts to identify regions of the genome some distance from functional sites or for which BGS is suggested to be absent (using the approach of McVicker *et al*. 2009, for example). Using diversity data from such regions, researchers should then estimate a demographic model. Care should be taken to simulate data under the estimated model to ensure it fits reasonably well with observations. Following the general trends outlined here (e.g., Figure 5) should then allow qualitative predictions about the impacts of demography and linked selection. We caution, though, that quantitative predictions will require simulations using a range of plausible DFEs using a genome structure relevant for the species of interest.

Finally, it is worth noting that our results are relevant not just for comparisons of *π* in regions of high and low effects of selection at linked sites, but also apply to comparisons of selected and neutral polymorphisms as well. Indeed, similar patterns have been observed for comparisons of selected and putatively neutral polymorphisms in both simulations and empirical data (Do *et al*. 2015; Koch and Novembre 2017; Simons and Sella 2016) and further demonstrate that differential dynamics of diversity in response to demography are ubiquitous across the genome.

## Conclusions

Genetic diversity across the genome is determined by the complex interplay of mutation, demographic history, and the effects of both direct and linked natural selection. While each of these processes is understood to a degree on its own, in many cases we lack either theory or sufficient empirical data to capture the effects of their interaction. Selection at linked sites, in particular, is increasingly recognized as perhaps the primary determinant of patterns of diversity along a chromosome (Comeron 2014; Stankowski *et al*. 2019), but our ability to infer its impact is often complicated by changes in population size. Many studies interested in these dynamics, however, make the simplifying assumption that selection at linked sites in such non-equilibrium populations can be effectively modeled using classic theory and scaling of the effective population size. Our extensive simulations show that, in the context of purifying selection, this is not the case. We find that the relationship between selection at linked sites and demographic change is complex, with shortterm dynamics often qualitatively different from predictions under classic models. These results suggest that inferring the impact of population size change on selection at linked sites should be undertaken with caution and is only really possible with a thorough understanding of the demographic history of the populations of interest.

## Acknowledgments

MGS and JR-I would like to acknowledge funding from NSF Plant Genome Grant 1238014 and funding from the Deutsche Forschungsgemeinschaft (DFG) grant STE 2654/1-1 to MGS. RDH was partially supported by grant R01HG007644 from the National Institutes of Health, and RT was supported by a Diversity Supplement under this award. We would also like to thank Felix Andrews for statistical advice, although we did not follow it.

---

**Figure S1.**
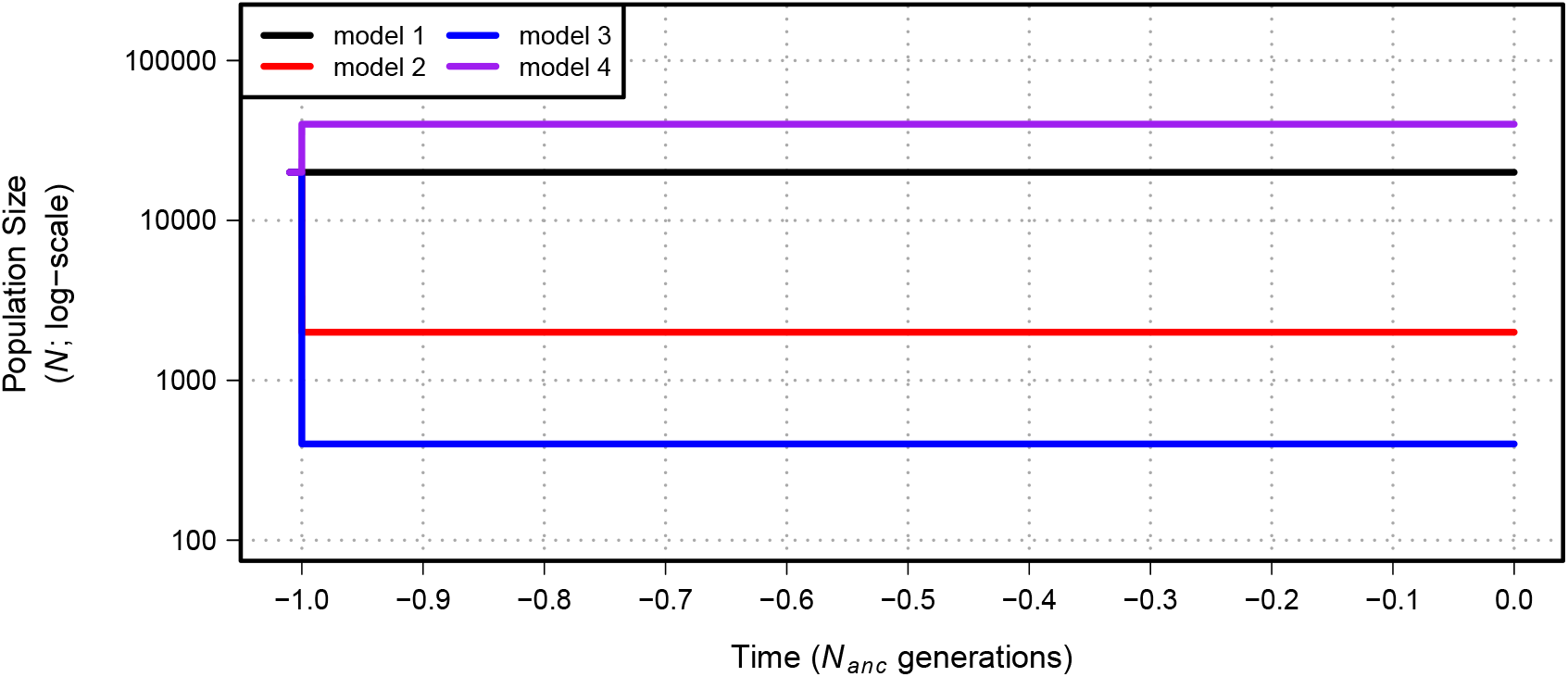
Demographic models 1-4 simulated in our study. Time proceeds forward from left to right and is scaled by the *N* of the population at the initial generation (*N_anc_*; 20,000 individuals). Demographic model 2 experiences a population contraction to 2000 individuals while demographic model 3 experiences a population contraction to 400 individuals. Demographic model 4 experiences a population expansion to 40,000 individuals. All population size changes are instantaneous for models 2-4. See Table 1 for additional model parameters.

**Figure S2.**
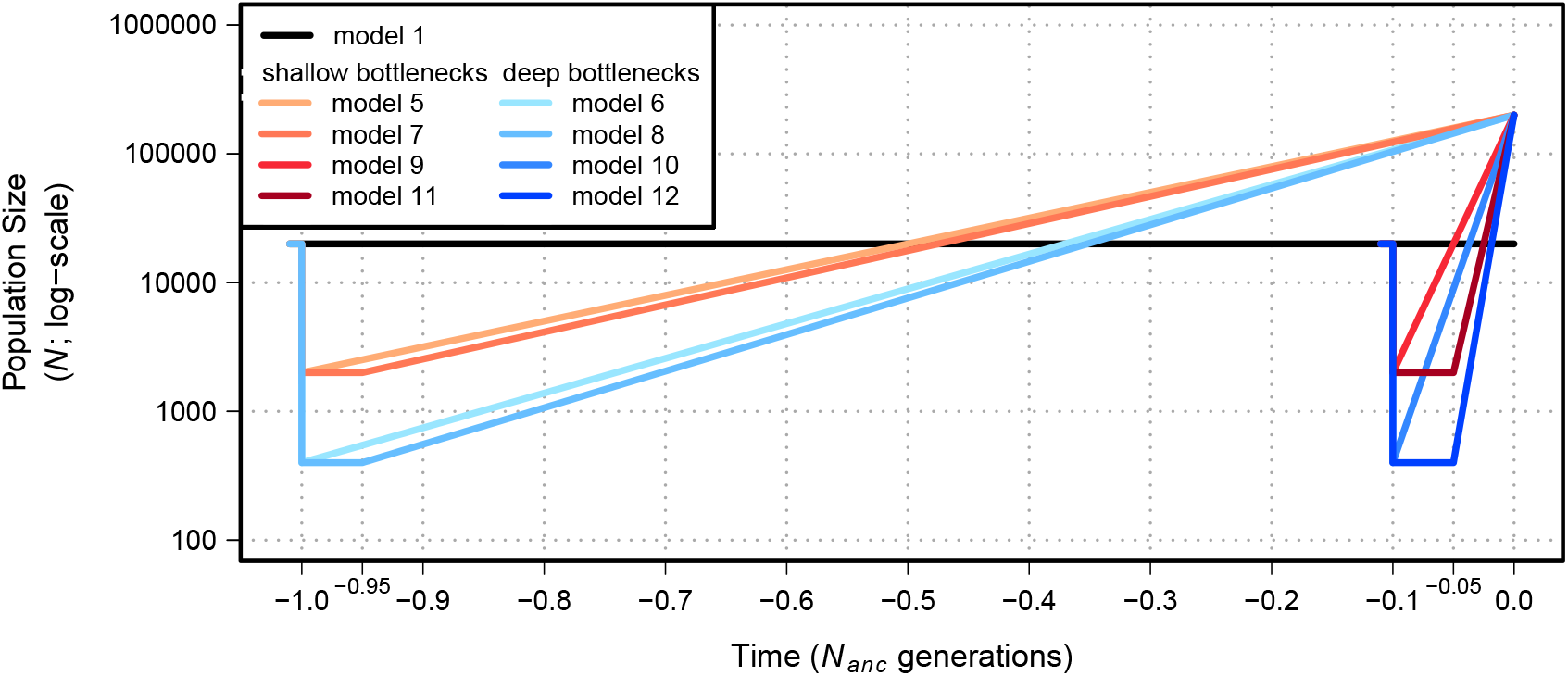
Demographic models 1 and 5-12 simulated in our study. Time proceeds forward from left to right and is scaled by the *N* of the population at the initial generation (*N_anc_*; 20,000 individuals). Demographic models with a shallow bottleneck (models 5, 7, 9, and 11) experience a population contraction to 2000 individuals while demographic models with a deep bottleneck (models 6, 8, 10, and 12) experience a population contraction to 400 individuals. After contraction, demographic models 5-12 undergo exponential growth to a final population size of 200,000 individuals. See Table 1 for additional model parameters.

**Figure S3.**
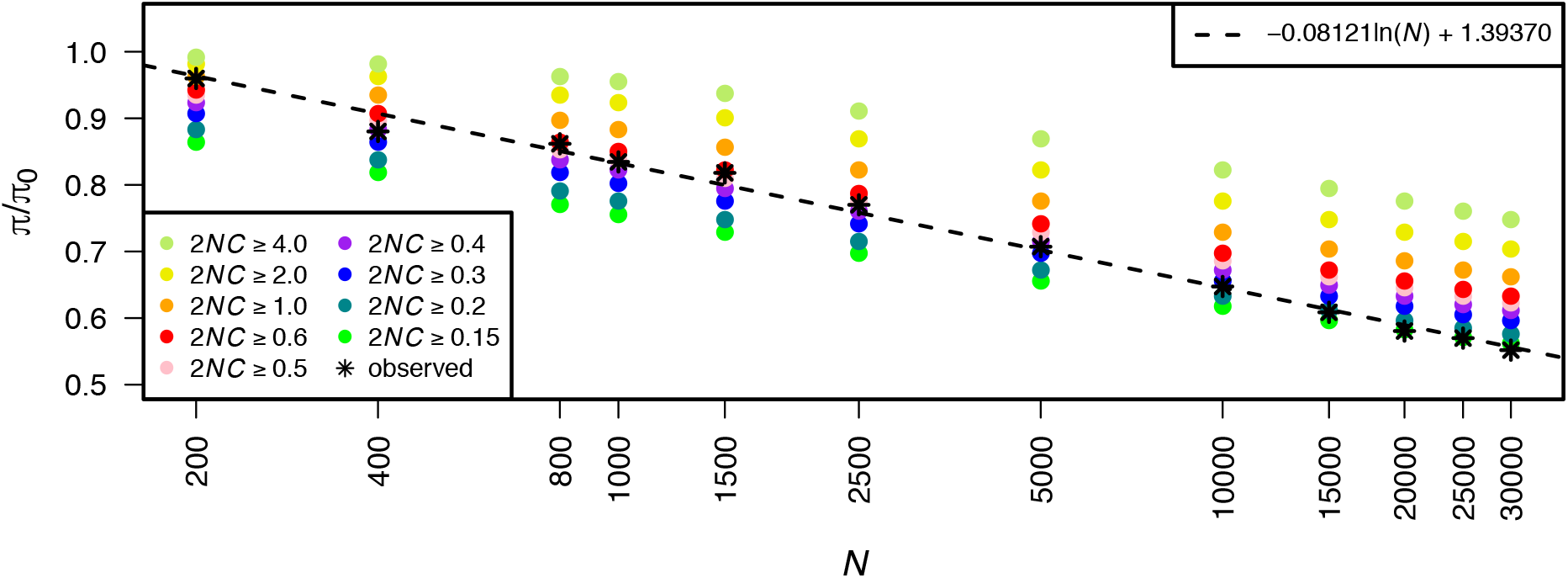
Estimate of *π*/*π*_0_ from the classic model (Nordborg *et al*. 1996) across different population sizes and different truncation thresholds on selection. Different *γ* values used to truncate selection (*C*) for the classic model are shown in the legend (2NC ≥ *γ*). Black stars represent the observed *π*/*π*_0_ from running simulations of BGS. The dashed line shows a log-linear model of *π*/*π*_0_ fit to the simulations of BGS (*π*/*π*_0_ = –0.08121ln(*N*) + 1.39370).

**Figure S4.**
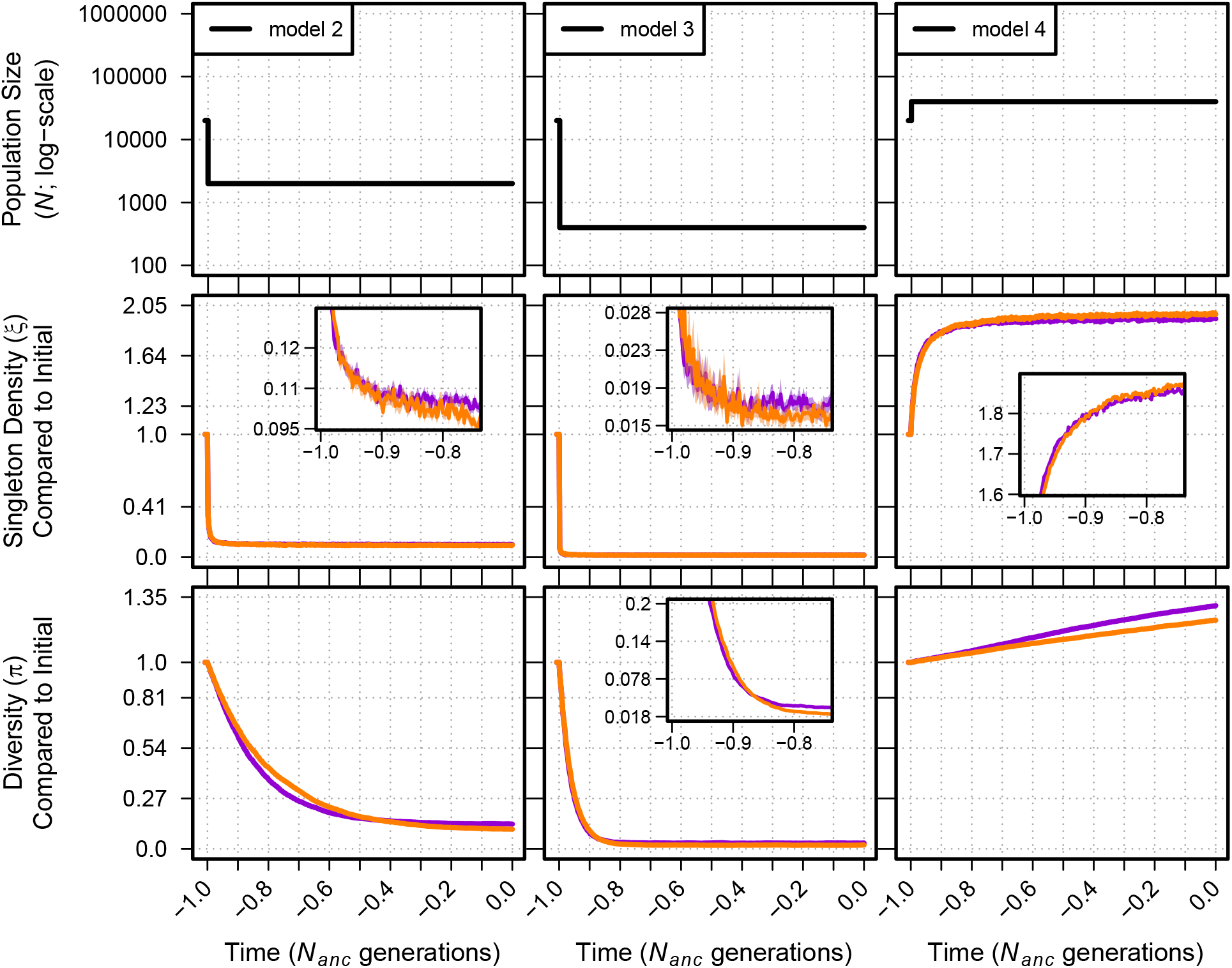
Singleton density (*ξ*) and diversity (*π*) for demographic models 2-4 under neutrality (orange lines) and BGS (violet lines) relative to their values in the initial generation prior to demographic change. The top panel shows each demographic model as in Figure 1. For greater detail, insets show data for generations over a smaller time scale and smaller y-axis (note: y-axes for insets are scaled linearly). Envelopes are 95% CIs calculated from 10,000 bootstraps of the original simulation data. The data used for this figure is identical to that of Figure 1.

**Figure S5.**
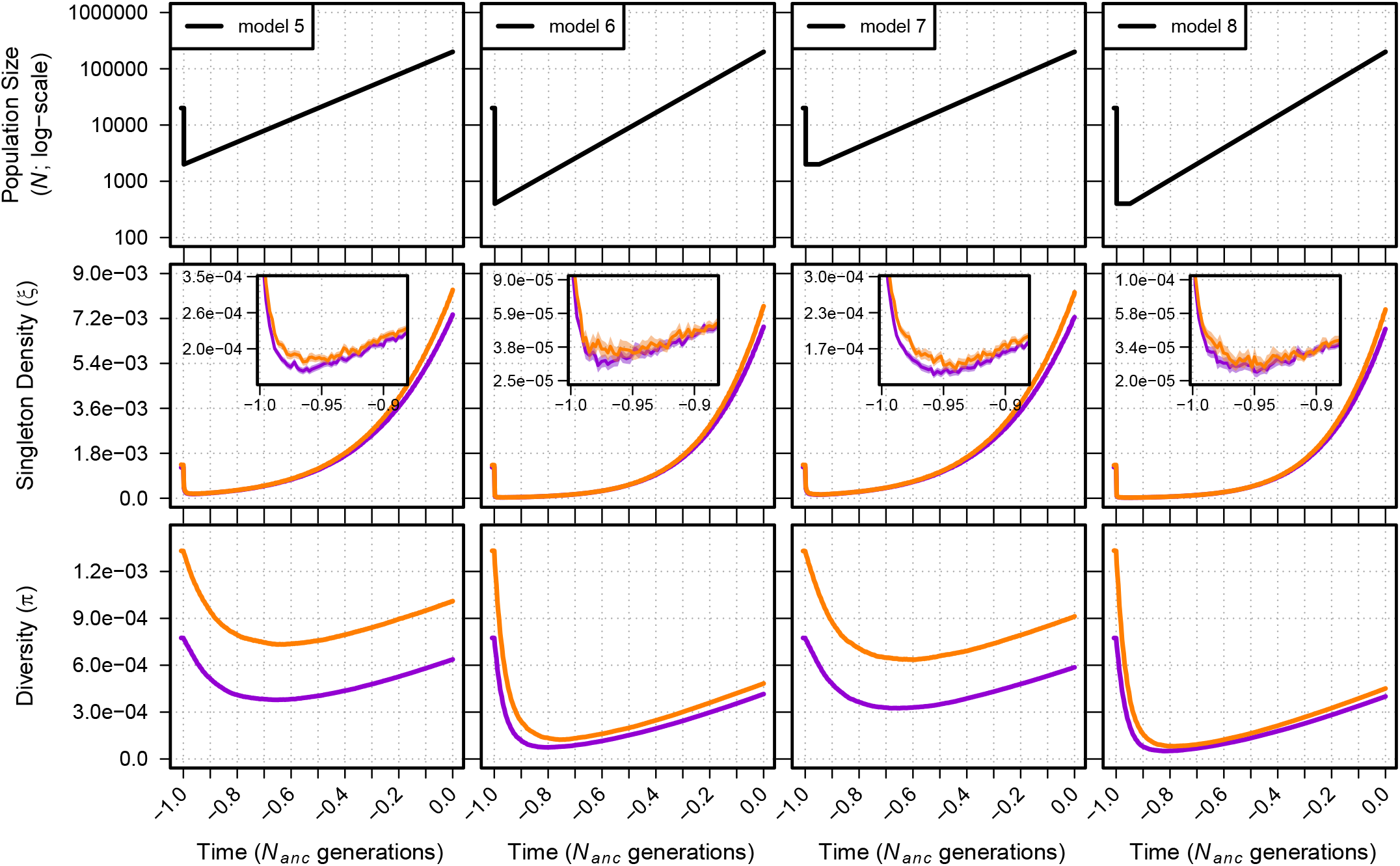
Singleton density (*ξ* per site) and diversity (*π* per site) for models 5-8. The top panel shows each demographic model; time proceeds forward from left to right and is scaled by the *N* of the population at the initial generation (*N_anc_*; 20,000 individuals). Diversity statistics are shown for neutral simulations (orange lines) and simulations with BGS (violet lines). Insets show diversity using a log scale for improved detail. Envelopes are 95% CIs calculated from 10,000 bootstraps of the original simulation data.

**Figure S6.**
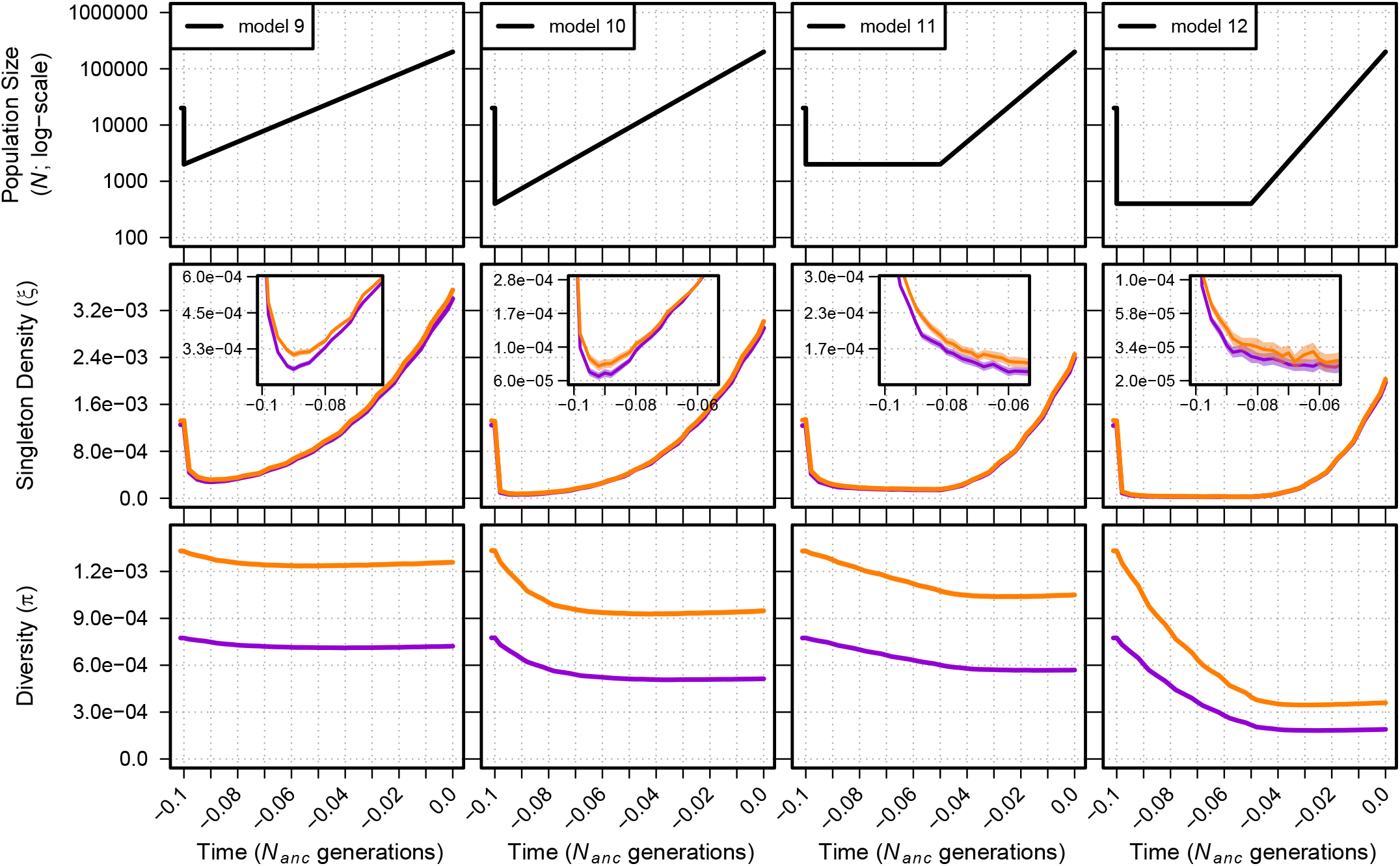
Singleton density (*ξ* per site) and diversity (*π* per site) for models 9-12. The top panel shows each demographic model; time proceeds forward from left to right and is scaled by the *N* of the population at the initial generation (*N_anc_*; 20,000 individuals). Diversity statistics are shown for neutral simulations (orange lines) and simulations with BGS (violet lines). Insets show diversity using a log scale for improved detail. Envelopes are 95% CIs calculated from 10,000 bootstraps of the original simulation data.

**Figure S7.**
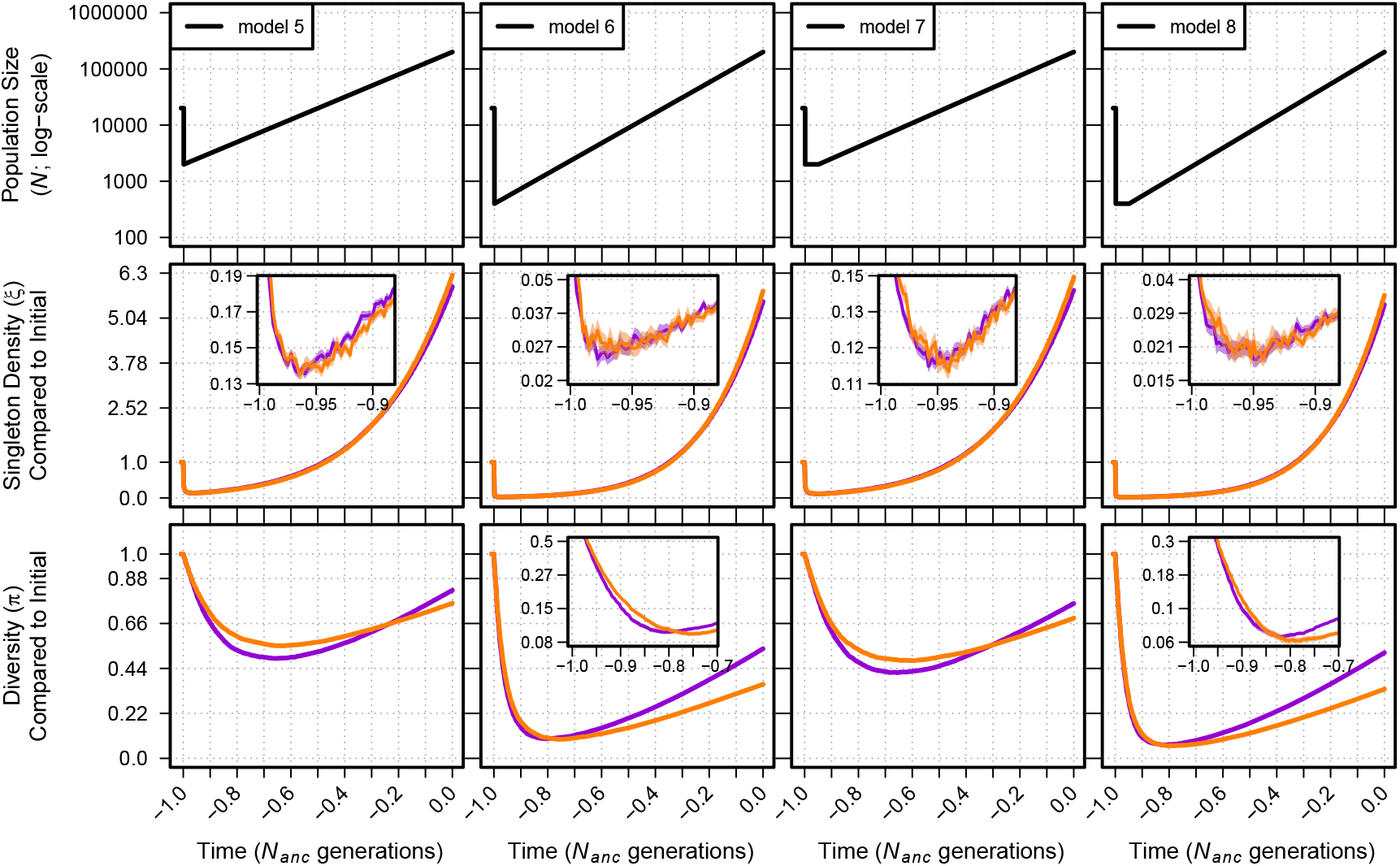
Singleton density (*ξ*) and diversity (*π*) relative to the initial generation for neutral (orange) and BGS (violet) simulations of demographic models 5-8. The top panel shows each demographic model as in Supplemental Figure S5. Insets show diversity over a shorter timescale and use a log scale for diversity for improved detail. Envelopes are 95% CIs calculated from 10,000 bootstraps of the original simulation data. The data used for this figure is identical to that of Supplemental Figure S5.

**Figure S8.**
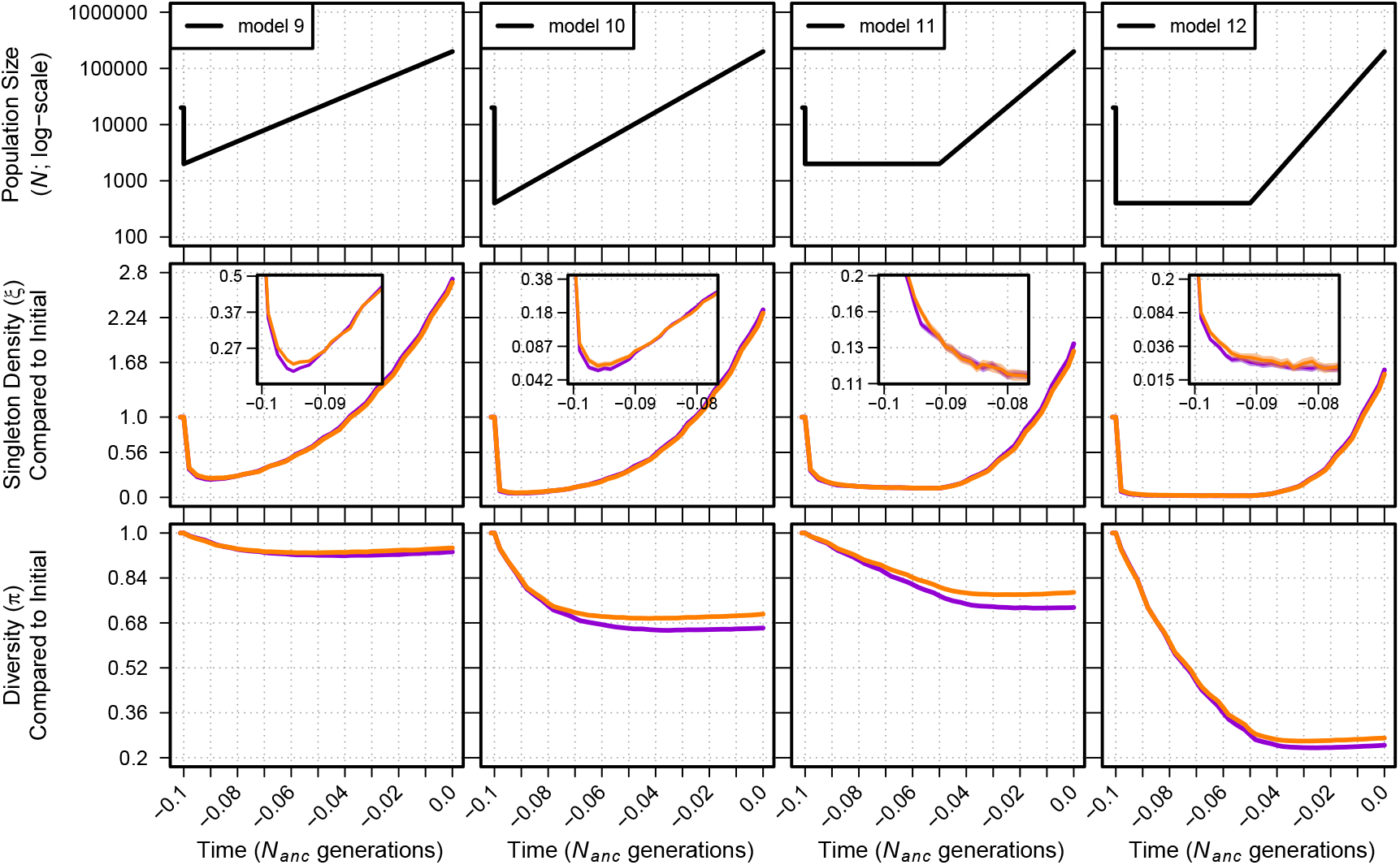
Singleton density (*ξ*) and diversity (*π*) relative to the initial generation for neutral (orange) and BGS (violet) simulations of demographic models 9-12. The top panel shows each demographic model as in Supplemental Figure S6. Insets show diversity over a shorter timescale and use a log scale for diversity for improved detail. Envelopes are 95% CIs calculated from 10,000 bootstraps of the original simulation data. The data used for this figure is identical to that of Supplemental Figure S6.

**Figure S9.**
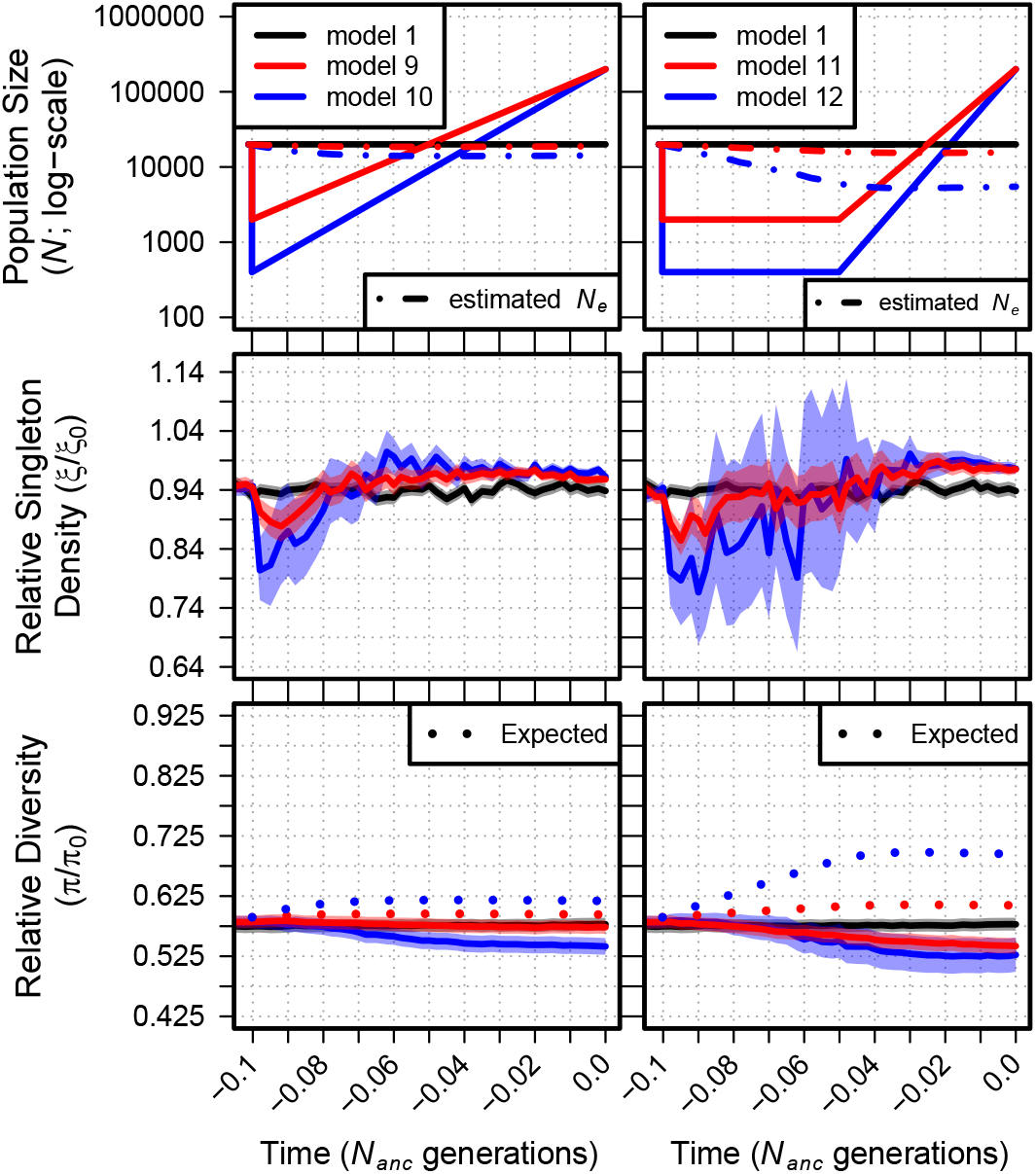
Relative singleton density (*ξ*/*ξ*_0_) and relative diversity (*π*/*π*_0_) across time for demographic models 1 and 9-12. The top panel shows each demographic model; time proceeds forward from left to right and is scaled by the *N* of the population at the initial generation (*N_anc_;* 20,000 individuals). Black lines show *ξ*/*ξ*_0_ and *π*/*π*_0_ from simulations of a constant sized population (model 1). Dot-dashed lines in the top panels show the estimated *N_e_* from observed *π*_0_. Dotted lines in the bottom panel show the equilibrium expectation of *π*/*π*_0_ from a log-linear model of simulated BGS with the specific selection parameters and the estimated *N_e_* at each time point (see Figure S3). Envelopes are 95% CIs calculated from 10,000 bootstraps of the original simulation data.

**Figure S10.**
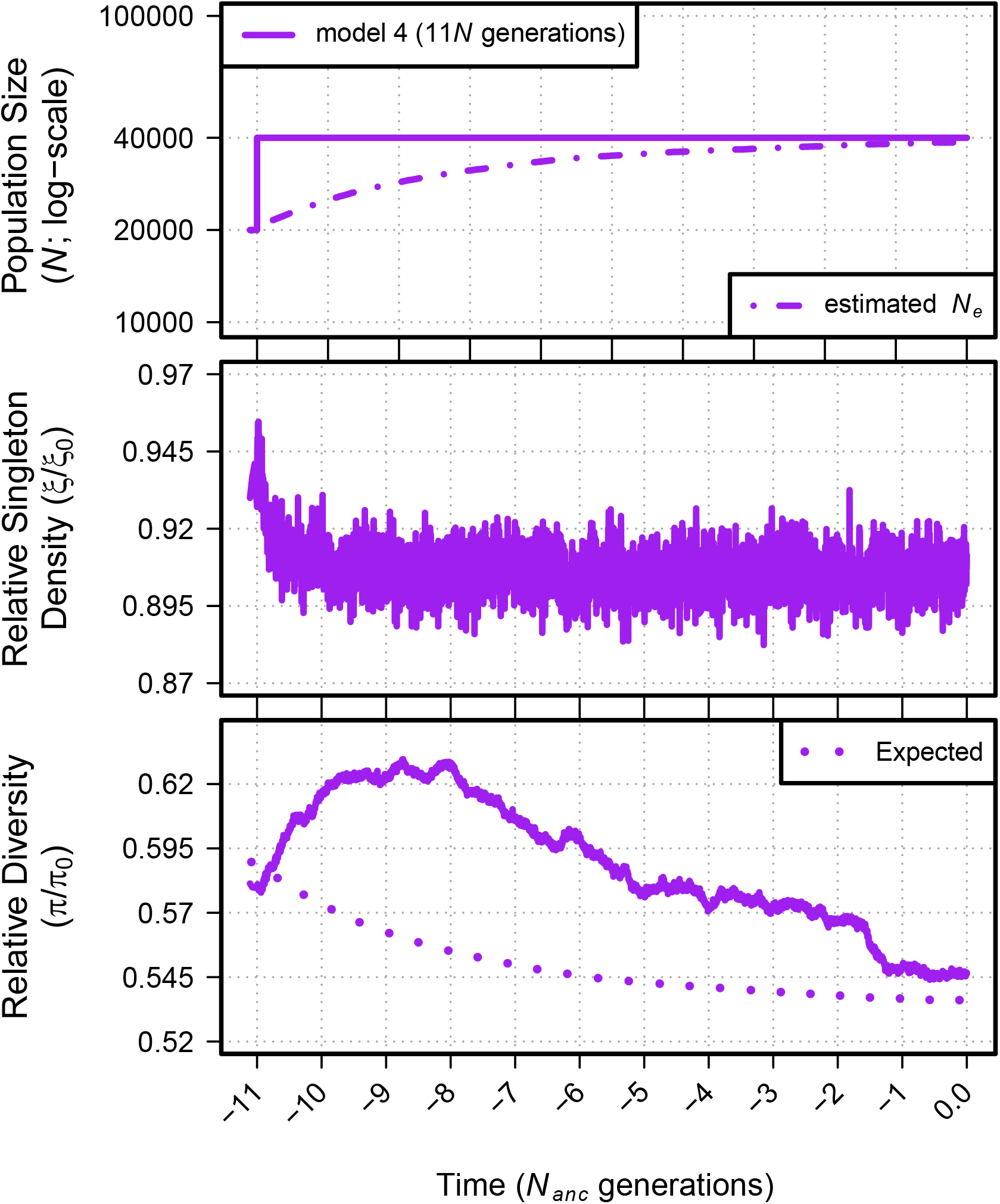
Relative singleton density (*ξ*/*ξ*_0_) and relative diversity (*π*/*π*_0_) across 11 *N_anc_* generations for demographic model 4. The top panel shows the demographic model; time proceeds forward from left to right and is scaled by the *N* of the population at the initial generation (*N_anc_;* 20,000 individuals). Dot-dashed lines in the top panels show the estimated *N_e_* from observed *π*_0_. Dotted lines in the bottom panel show the equilibrium expectation of *π*/*π*_0_ from a log-linear model of simulated BGS with the specific selection parameters and the estimated *N_e_* at each time point (see Figure S3).

**Figure S11.**
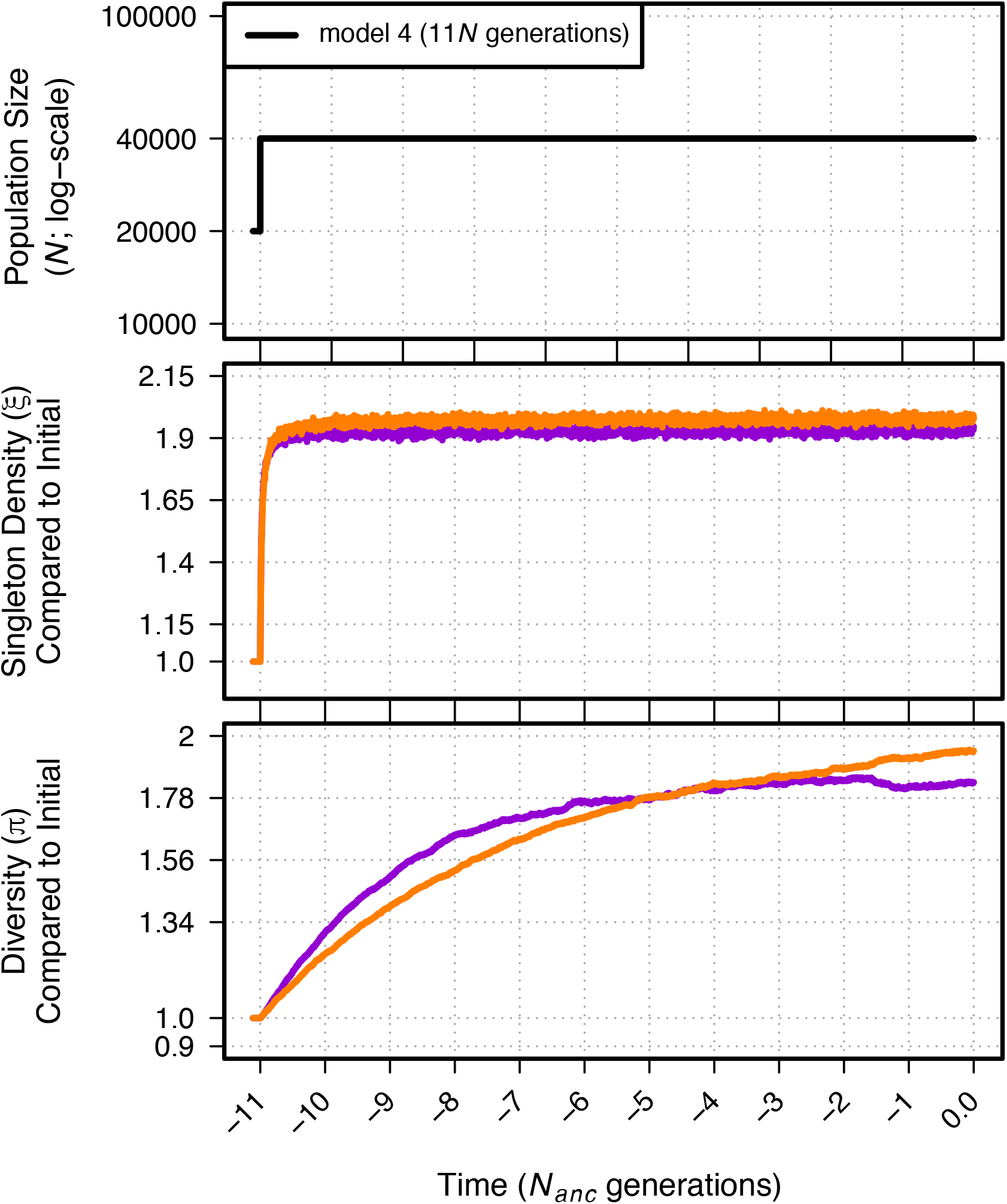
Singleton density (*ξ*) and diversity (*π*) relative to the initial generation for neutral (orange) and BGS (violet) simulations of demographic model 4 over 11 *N_anc_* generations. The top panel shows the demographic model.

**Figure S12.**
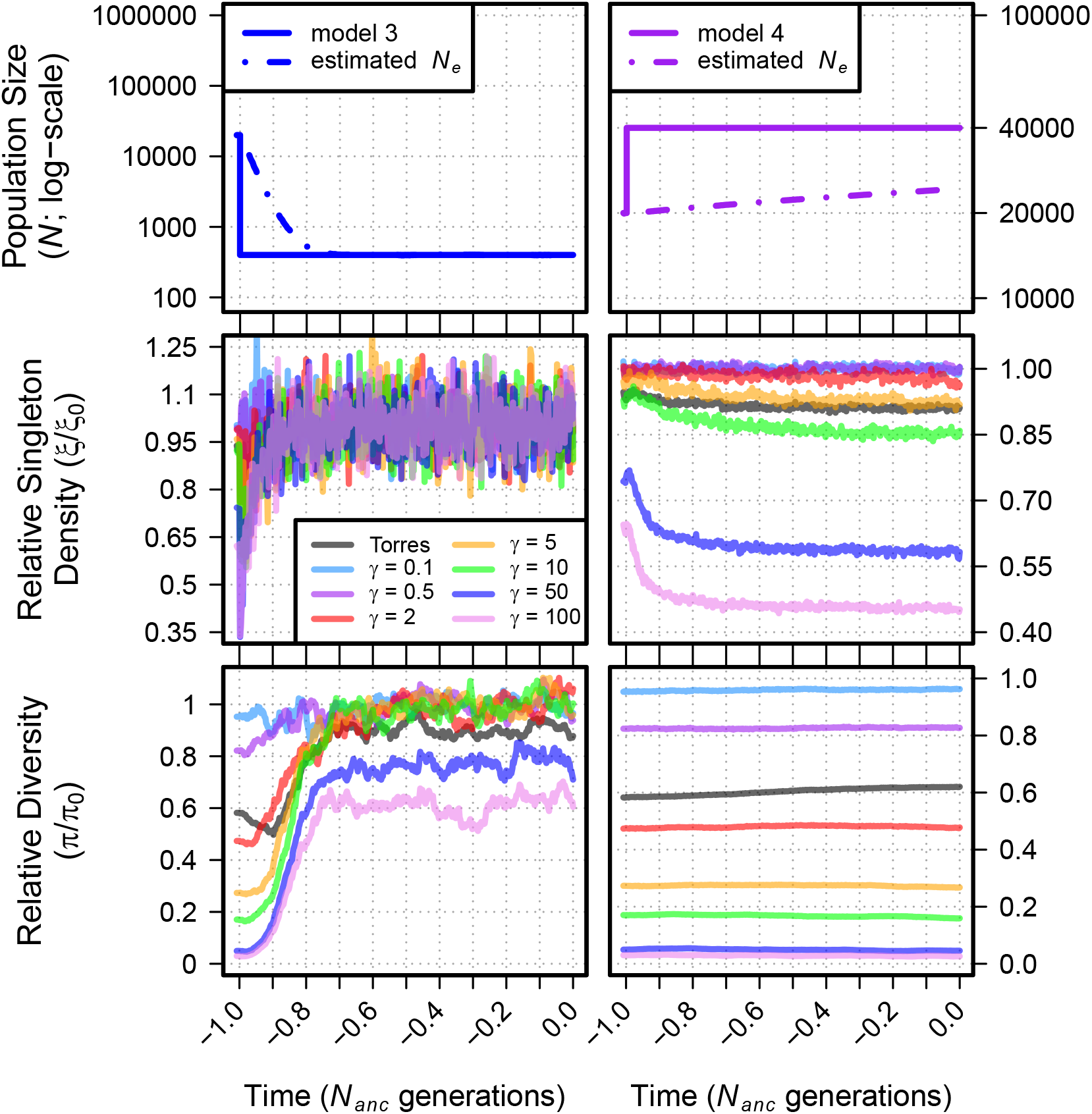
Relative singleton density (*ξ*/*ξ*_0_) and relative diversity (*π*/*π*_0_) across time for demographic models 3 and 4 for varying values of *γ* (where *γ* = *2N_anc_s*). *γ* is drawn from a single value for these simulations such that mean and median *γ* are the same. For comparison, we include the results of our simulated DFE that was used for demographic models 1-12 (indicated as “Torres” in the legend; mean *γ* ≈ 424, median *γ* ≈ 0.056). The top panel shows the demographic model; time proceeds forward from left to right and is scaled by the *N* of the population at the initial generation (*N_anc_*; 20,000 individuals). Dot-dashed lines in the top panels show the estimated *N_e_* from observed *π*_0_. Lines for *π*/*π*_0_ and *ξ*/*ξ*_0_ are transparent to show greater detail.

**Figure S13.**
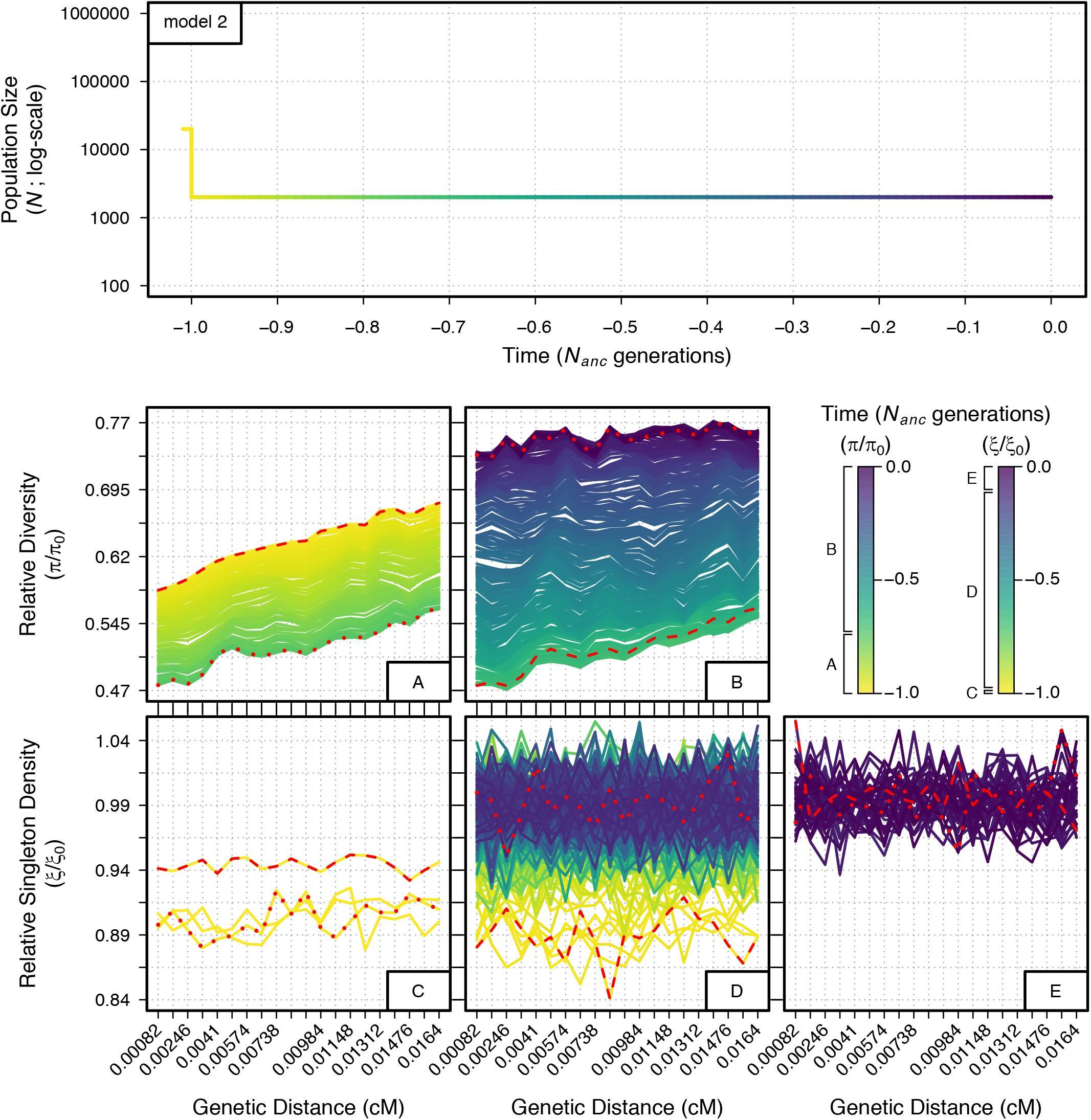
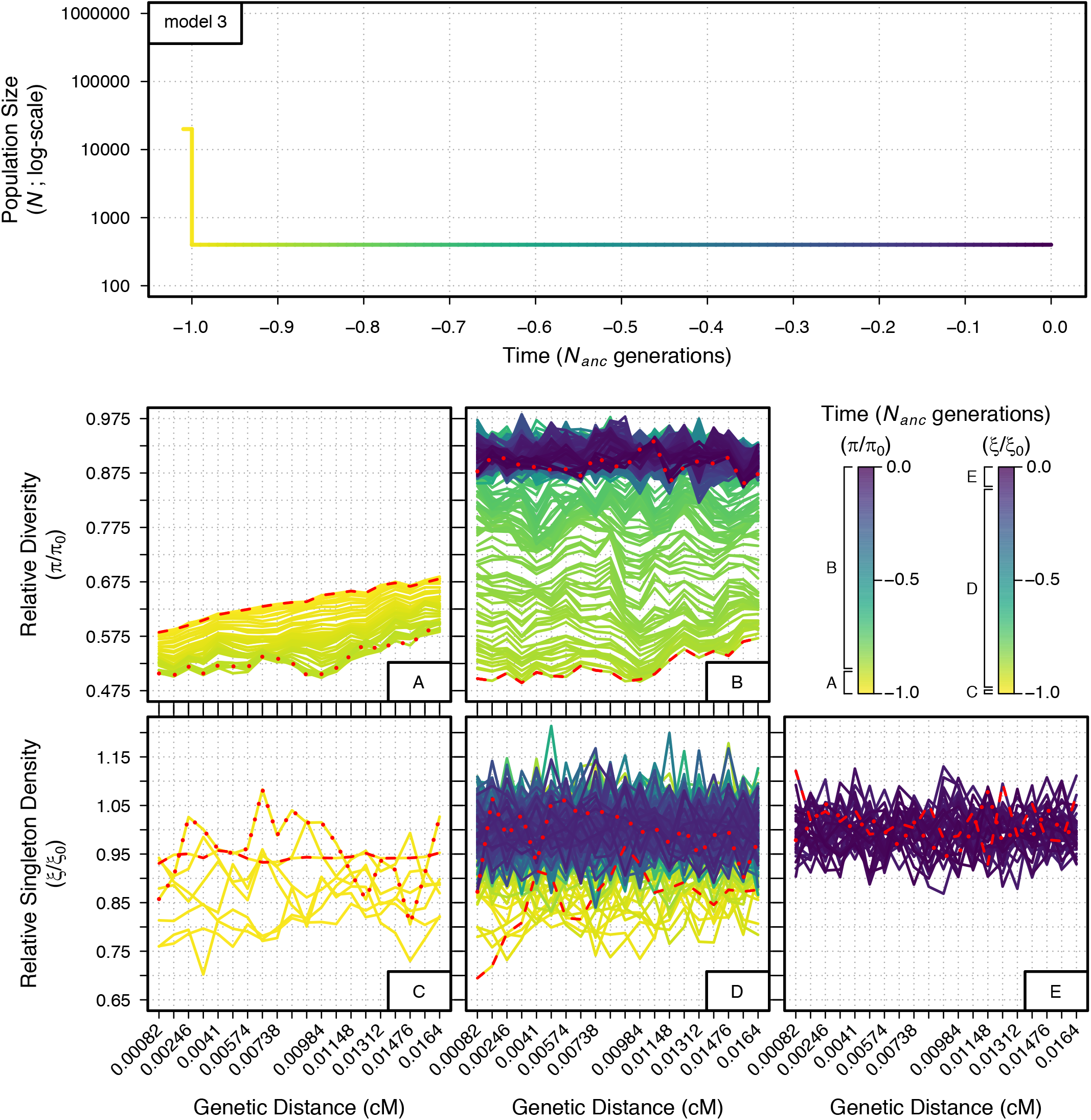
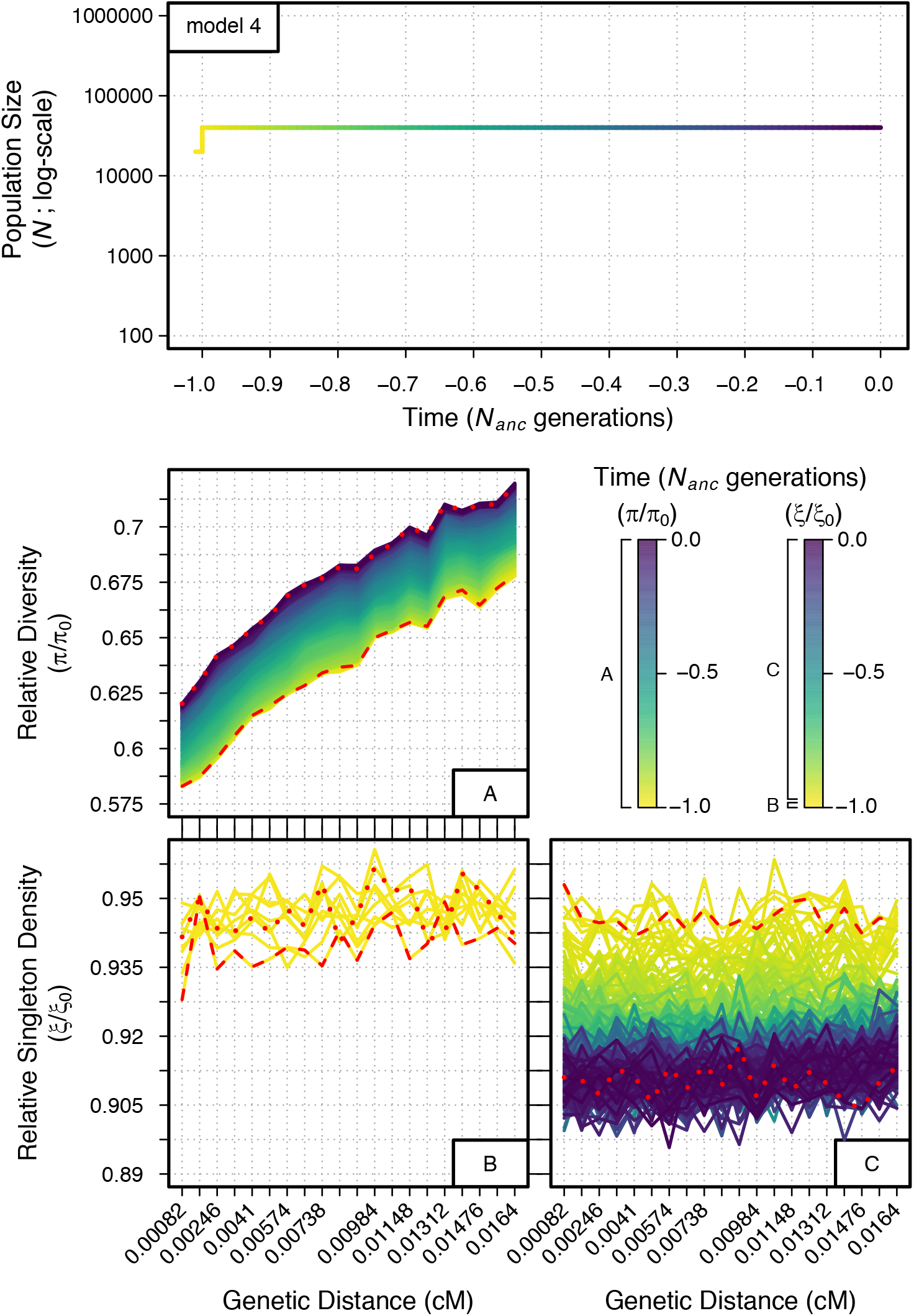
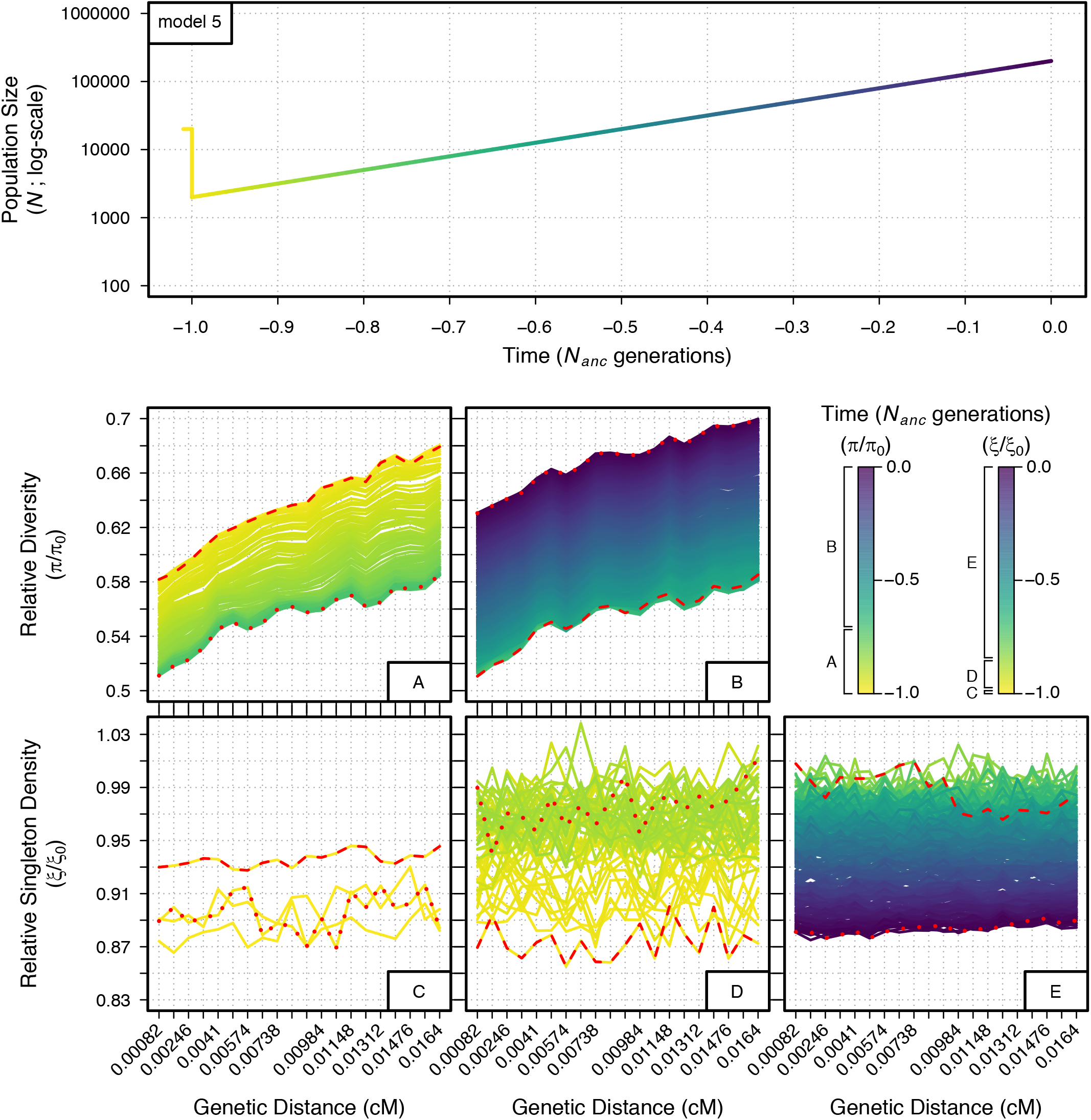
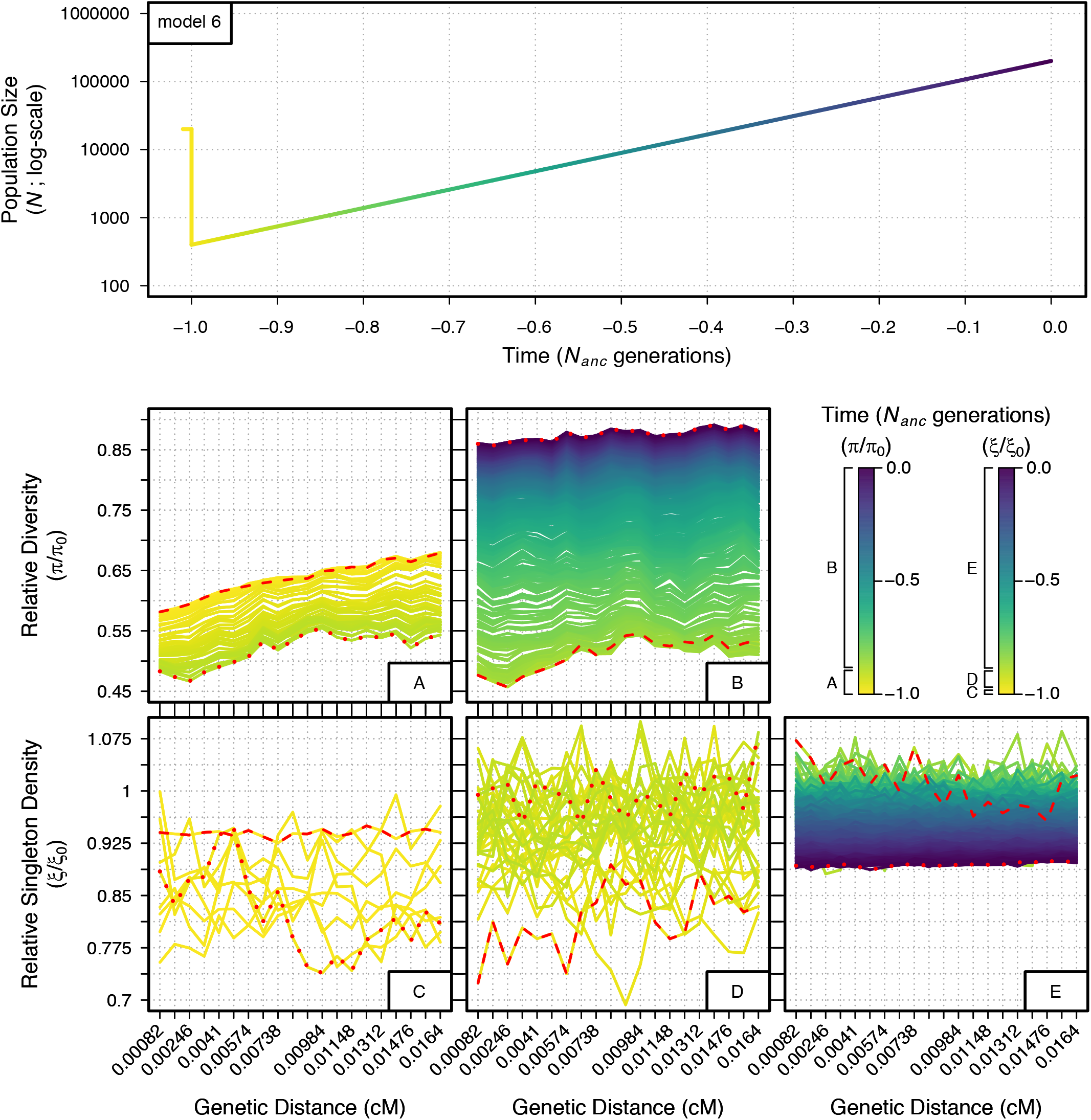
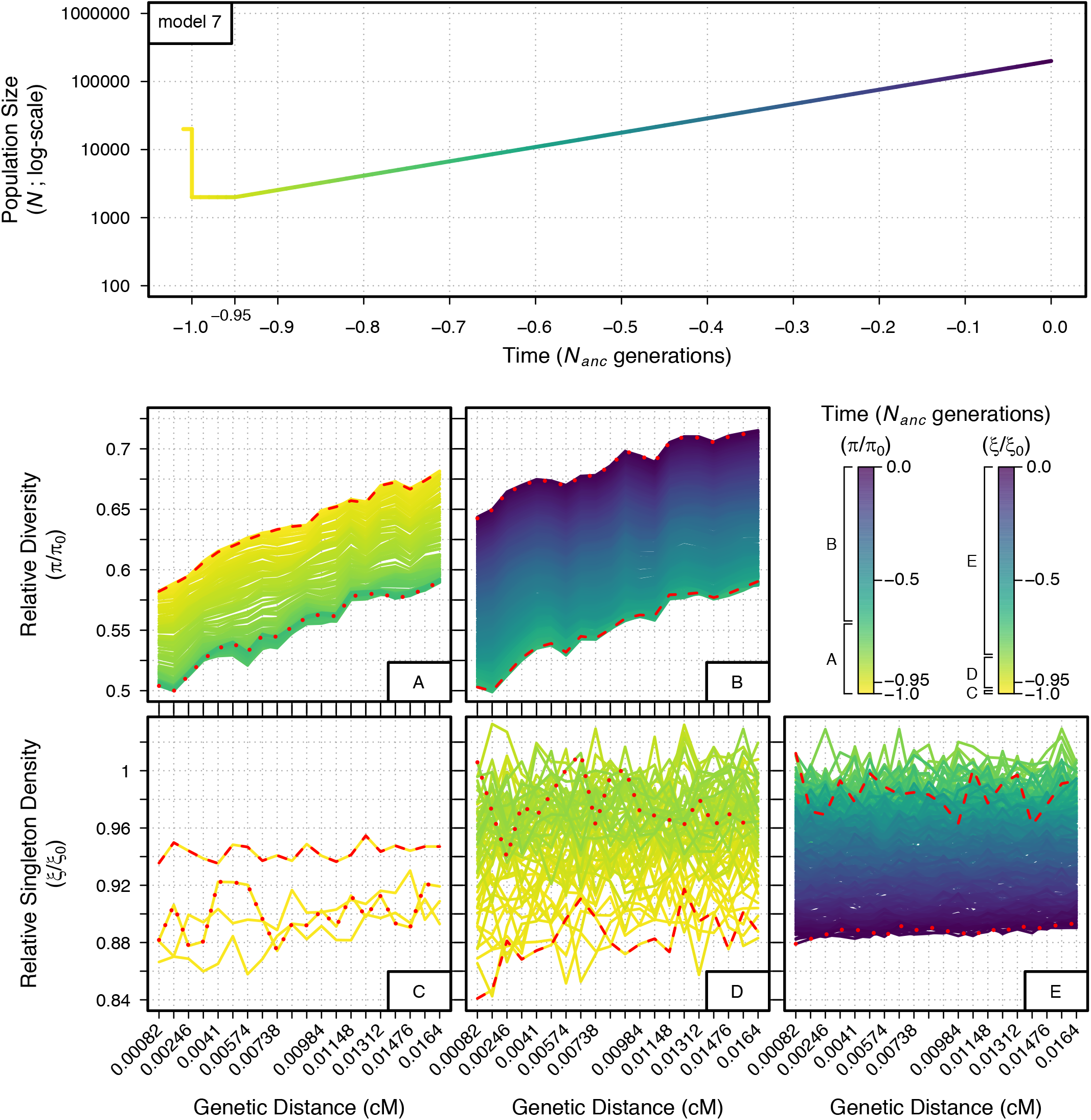
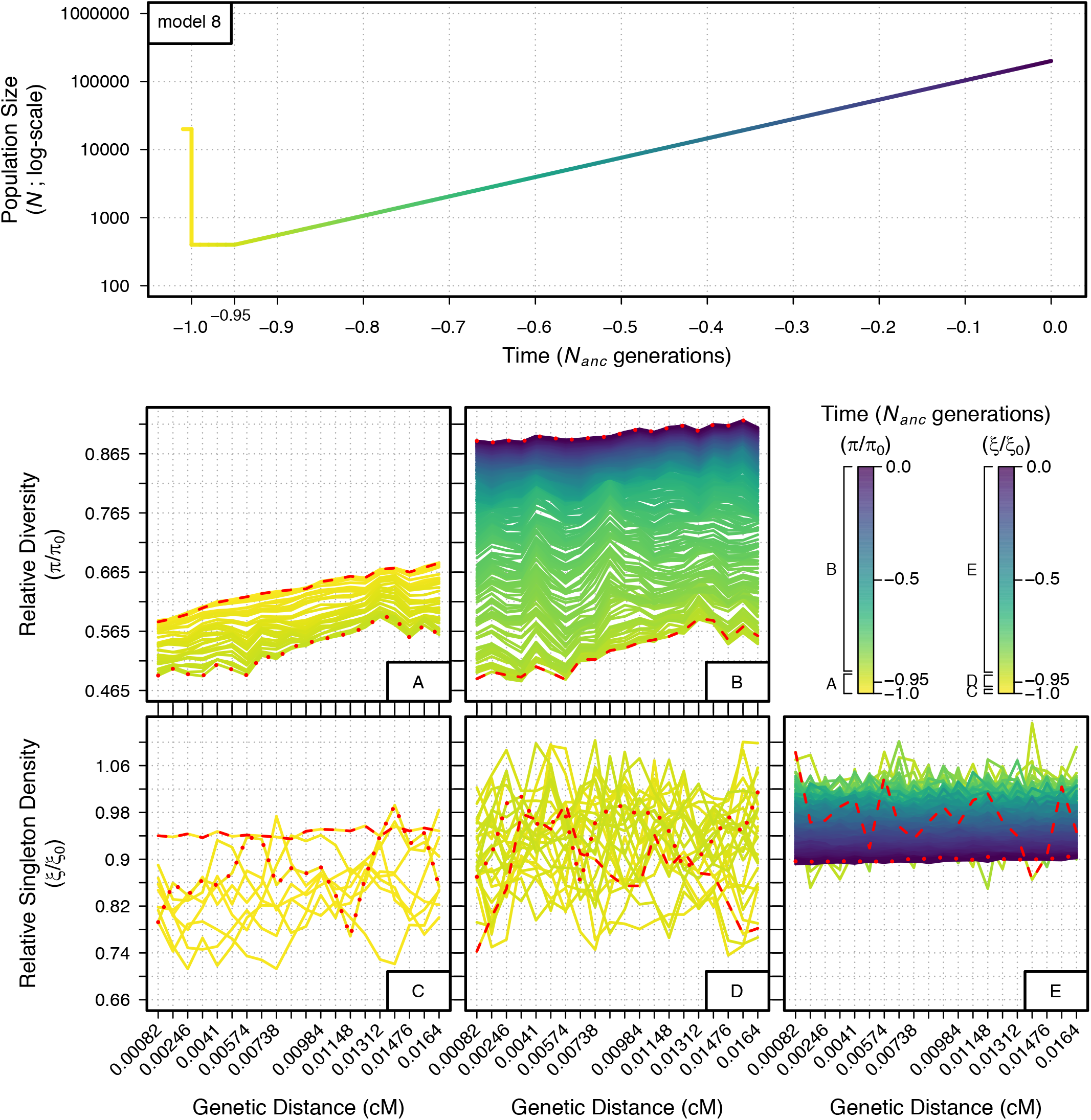
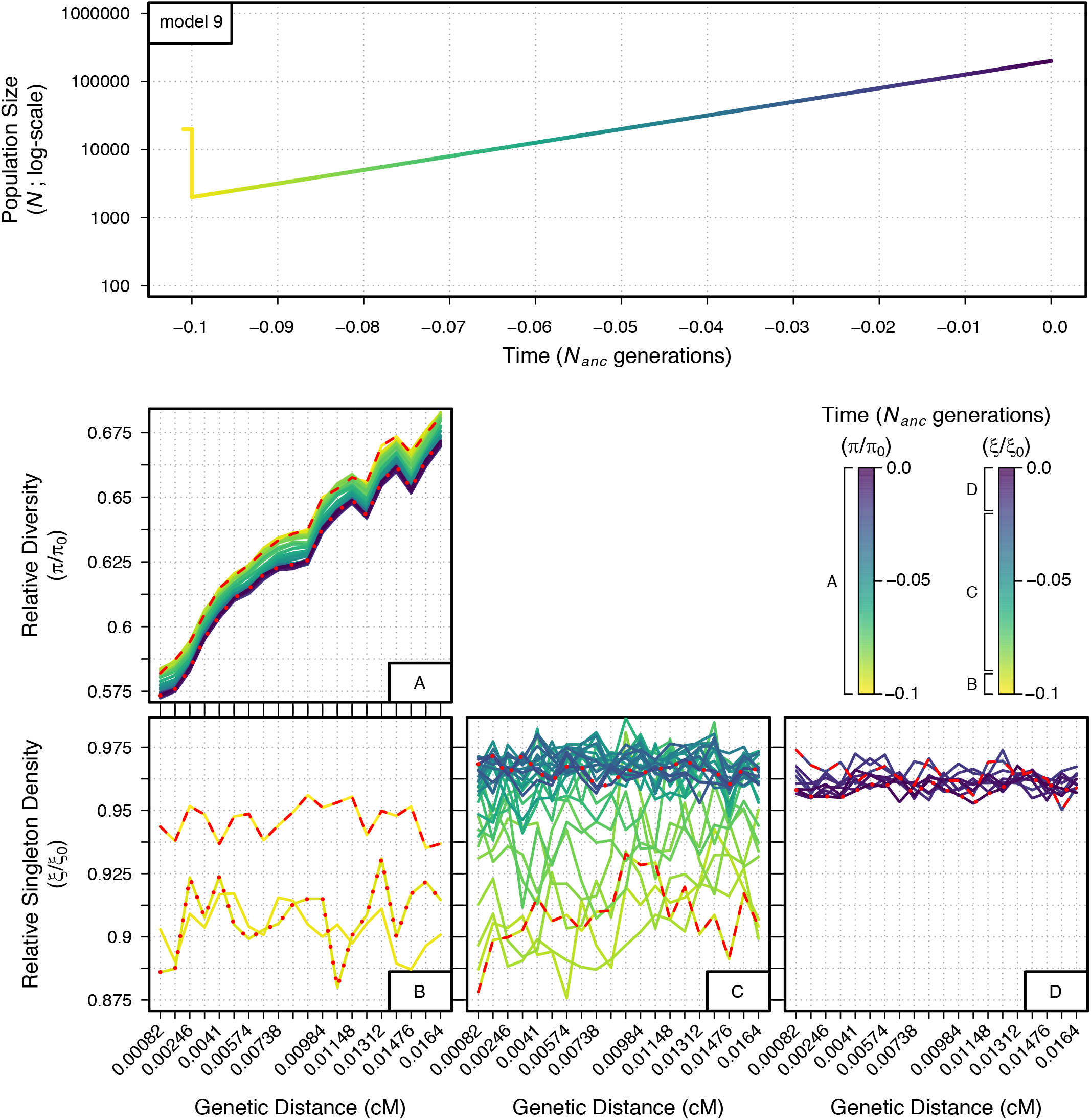
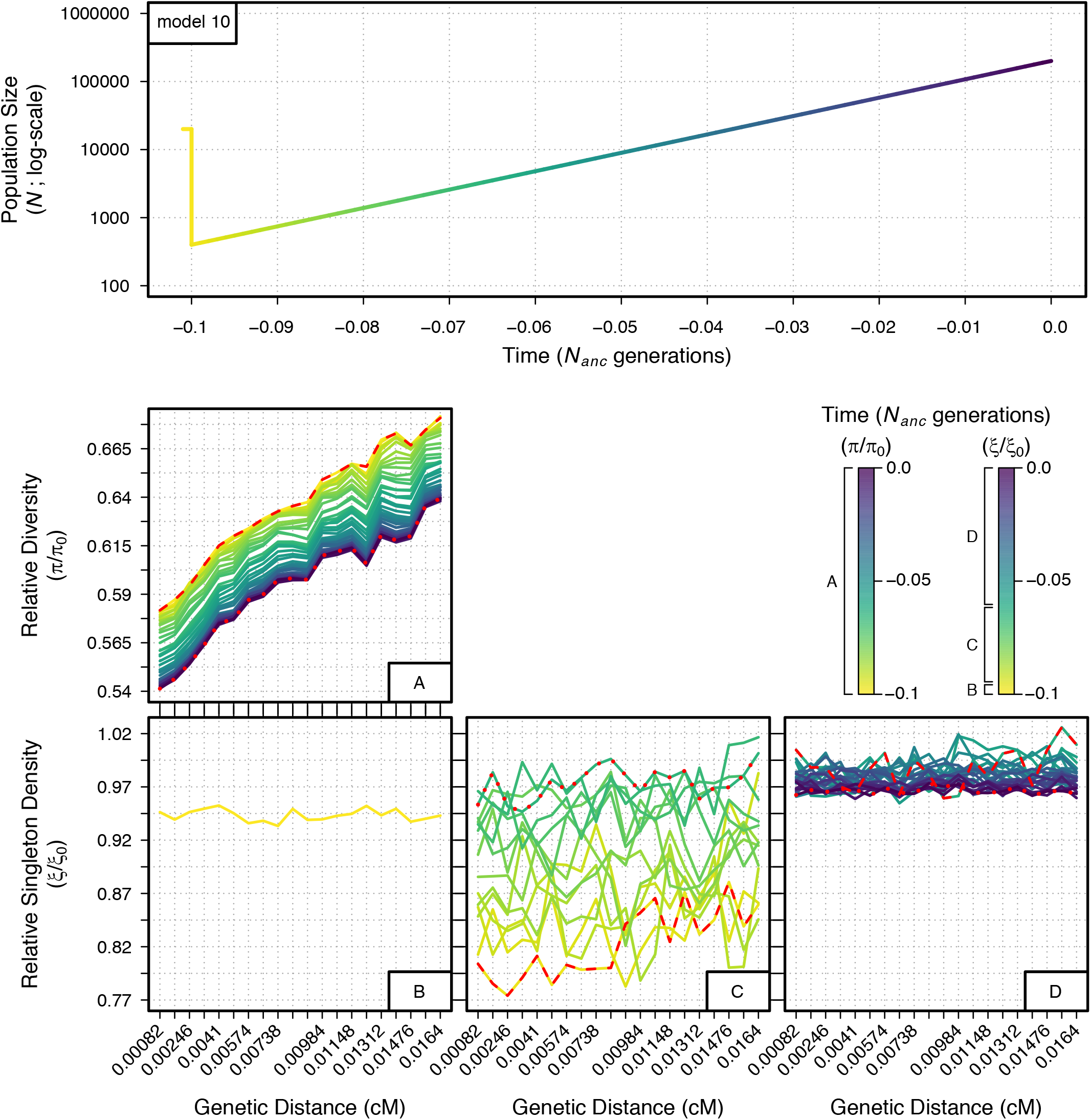
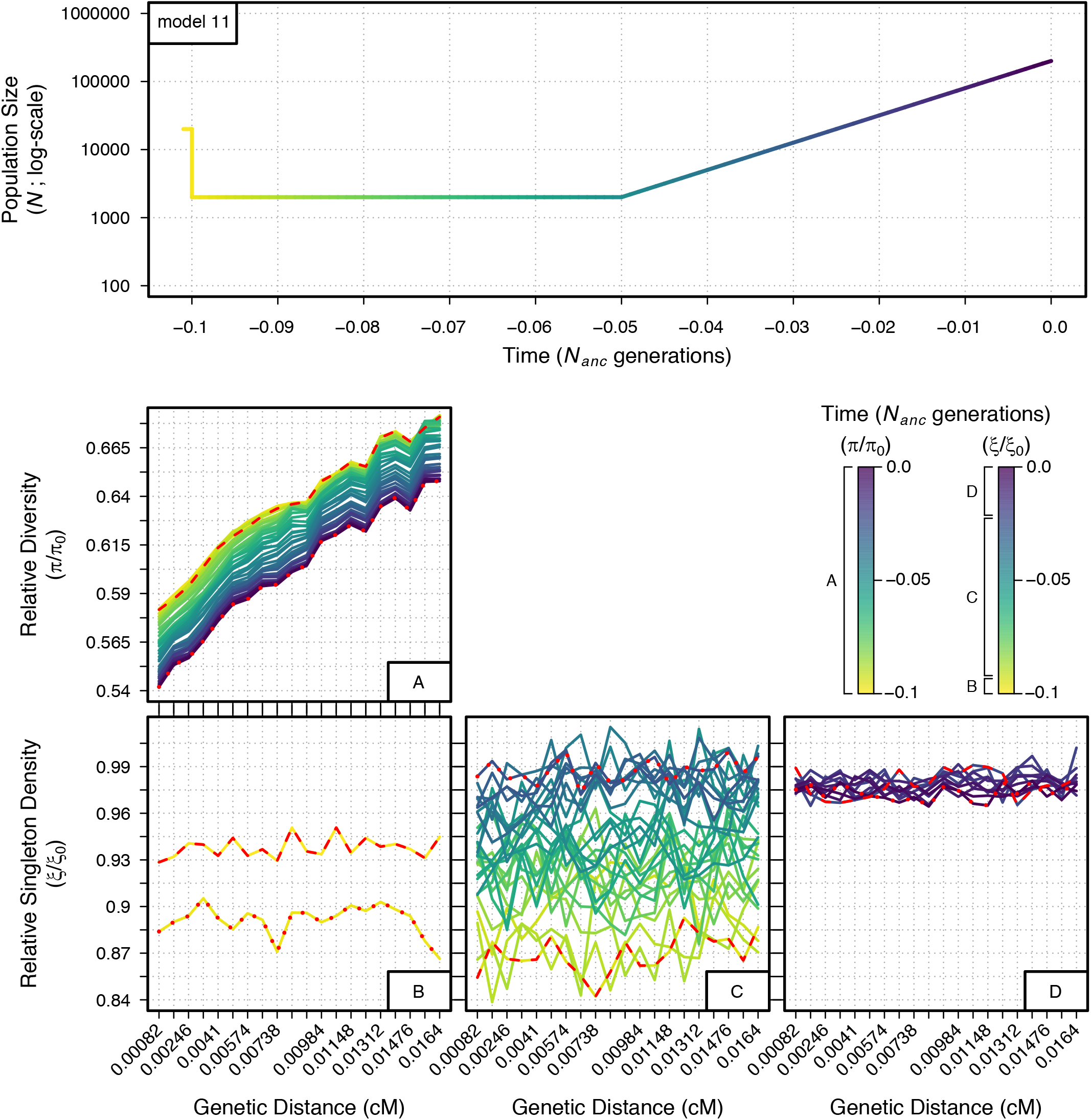
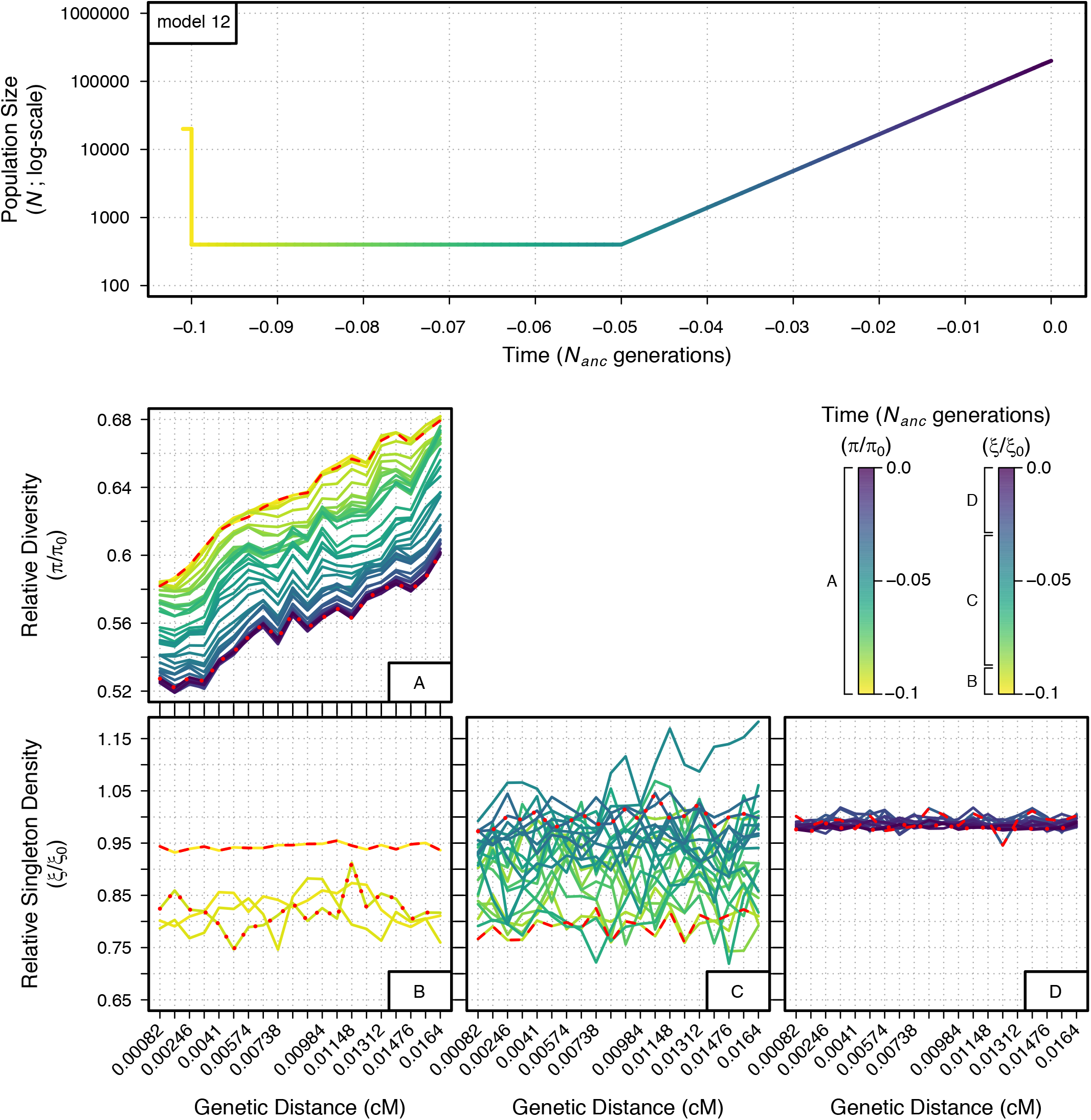
Relative diversity (*π*/*π*_0_) and singleton density (*ξ*/*ξ*_0_) through time for demographic models 2-12 measured across a neutral 200 kb region under the effects of BGS. The genetic distance of each 10 kb bin from the selected locus is indicated on the x-axes of the bottom two panels, with genetic distance increasing from left to right. Each line measuring *π*/*π*_0_ and *ξ*/*ξ*_0_ across the 200 kb neutral region represents a specific generation of the demographic model (401 discrete generations for demographic models 2-8, 41 discrete generations for demographic models 9-12). Specific generations are indicated by the color of the demographic model at the top of each figure (time is scaled in units of *N_anc_* generations [20,000 indiaryingiduals]) and in the figure legend. When necessary, multiple plots are given for *π*/*π*_0_ and *ξ*/*ξ*_0_ in order to prevent overlap of the measurements between generations (see legend for specific generations covered in each plot). Red dashed lines and red dotted lines indicate the first generation and last generation measured, respectively, for each specific plot.

